# Tracking the genetic diversity of SARS-CoV-2 variants in Nicaragua throughout the COVID-19 Pandemic

**DOI:** 10.1101/2024.06.03.596876

**Authors:** Gerald Vásquez Alemán, Cristhiam Cerpas, Jose G. Juarez, Hanny Moreira, Sonia Arguello, Josefina Coloma, Eva Harris, Aubree Gordon, Shannon N. Bennett, Ángel Balmaseda

## Abstract

The global circulation of SARS-CoV-2 has been extensively documented, yet the dynamics within Central America, particularly Nicaragua, remain underexplored. This study characterizes the genomic diversity of SARS-CoV-2 in Nicaragua from March 2020 through December 2022, utilizing 1064 genomes obtained via next-generation sequencing. These sequences were selected nationwide and analyzed for variant classification, lineage predominance, and phylogenetic diversity. We employed both Illumina and Oxford Nanopore Technologies for all sequencing procedures. Results indicated a temporal and spatial shift in dominant lineages, initially from B.1 and A.2 in early 2020 to various Omicron subvariants towards the study’s end. Significant lineage shifts correlated with changes in COVID-19 positivity rates, underscoring the epidemiological impact of variant dissemination. The comparative analysis with regional data underscored the low diversity of circulating lineages in Nicaragua and their delayed introduction compared to other countries in the Central American region. The study also linked specific viral mutations with hospitalization rates, emphasizing the clinical relevance of genomic surveillance. This research advances the understanding of SARS-CoV-2 evolution in Nicaragua and provide valuable information regarding its genetic diversity for public health officials in Central America. We highlight the critical role of ongoing genomic surveillance in identifying emergent lineages and informing public health strategies.

## INTRODUCTION

Severe Acute Respiratory Syndrome Coronavirus 2 (SARS-CoV-2) caused the most recent global pandemic of Coronavirus Disease 2019 (COVID-19)^1^. As of January 2023, 67 million cases have been reported worldwide, with the death toll surpassing 6.8 million^2^. Due to the novelty and severity of the virus, a complete genome was rapidly generated by January 2020 and published in February 2020^3^. The publication of this genome made possible the development of vaccines and subsequent global vaccination effort. While disease severity is thus greatly mitigated, SARS-CoV-2 continues to circulate and its evolutionary dynamics include mutations associated with improving replication, increased virulence, and/or changes in the clinical progression of patients^4–6^. New strategies for genomic surveillance through virus RNA sequencing have been implemented globally to meet this ongoing challenge^7^, making genomic surveillance accessible and indispensable for epidemiological research to understand the impact of viral evolution on COVID-19 patients.

Since emergence, SARS-CoV-2 has continued to evolve into a variety of lineages, often with different properties, that may or may not persist (e.g., ^8,9^). Sequencing technologies have been important in tracking these genetic changes and the emergence of new lineages. Indeed, in response to the COVID-19 pandemic, many countries have invested in genomic sequencing as a component of health care infrastructure to track specific lineages or variants of concern and gain a deeper understanding of the pandemic’s dynamics locally. For example, the introduction of new genetic variants into a population was associated with new waves of infection^10^. Variants differ in terms of transmission dynamics, disease severity, and mortality risk to infected individuals^11^. Finally, the introduction of new lineages has been associated with changes in the risk of infection and severity in different age groups^12,13^.

In Nicaragua, the COVID-19 pandemic began in mid-March 2020. Since then, several published studies have investigated the immunological impact of SARS-CoV-2 infection in various Nicaraguan cohorts^14–16^. However, the origin, diversity and dynamics of circulating lineages in Nicaragua have not been well-studied. Our study documents nation-wide transmission dynamics of SARS-CoV-2 lineages. Thus, this research also contributes insights into the evolution of SARS-CoV-2 in Central America where exists a lack of information regarding genomic sequences due to limited infrastructure and low investment on genomic surveillance.

## RESULTS

### SARS-CoV-2 genomes and associated data

Through epidemiological surveillance by the Ministry of Health in Nicaragua, nasopharyngeal swabs positive for SARS-CoV-2 via qRT-PCR with a Ct range of 18-30 were randomly selected for sequencing. A total of 1064 SARS-CoV-2 genomes with coverage exceeding 60% from all departments and autonomous regions of Nicaragua were recovered using Next Generation Sequencing with Illumina technology or Oxford Nanopore Technology (ONT) (Supplementary Fig. 1c), and included in this study. Of the genomes analyzed, 1062 (99.8%) had patient sex data, with 620 (58.3%) being female and 442 (41.5%) males. Additionally, hospitalization data were retrieved for 639 (60.1%) sequences (Supplementary Table 1). Over forty percent of the recovered genomes originate from the capital of the country, Managua, where the National Virology lab is located, followed by departments in the central region, Estelí and Matagalpa (7.89% and 7.33% respectively), and then the department of León (7.05%) on Nicaragua’s Pacific coast (Fig.2 and Supplementary table 3).

**Figure 1.**
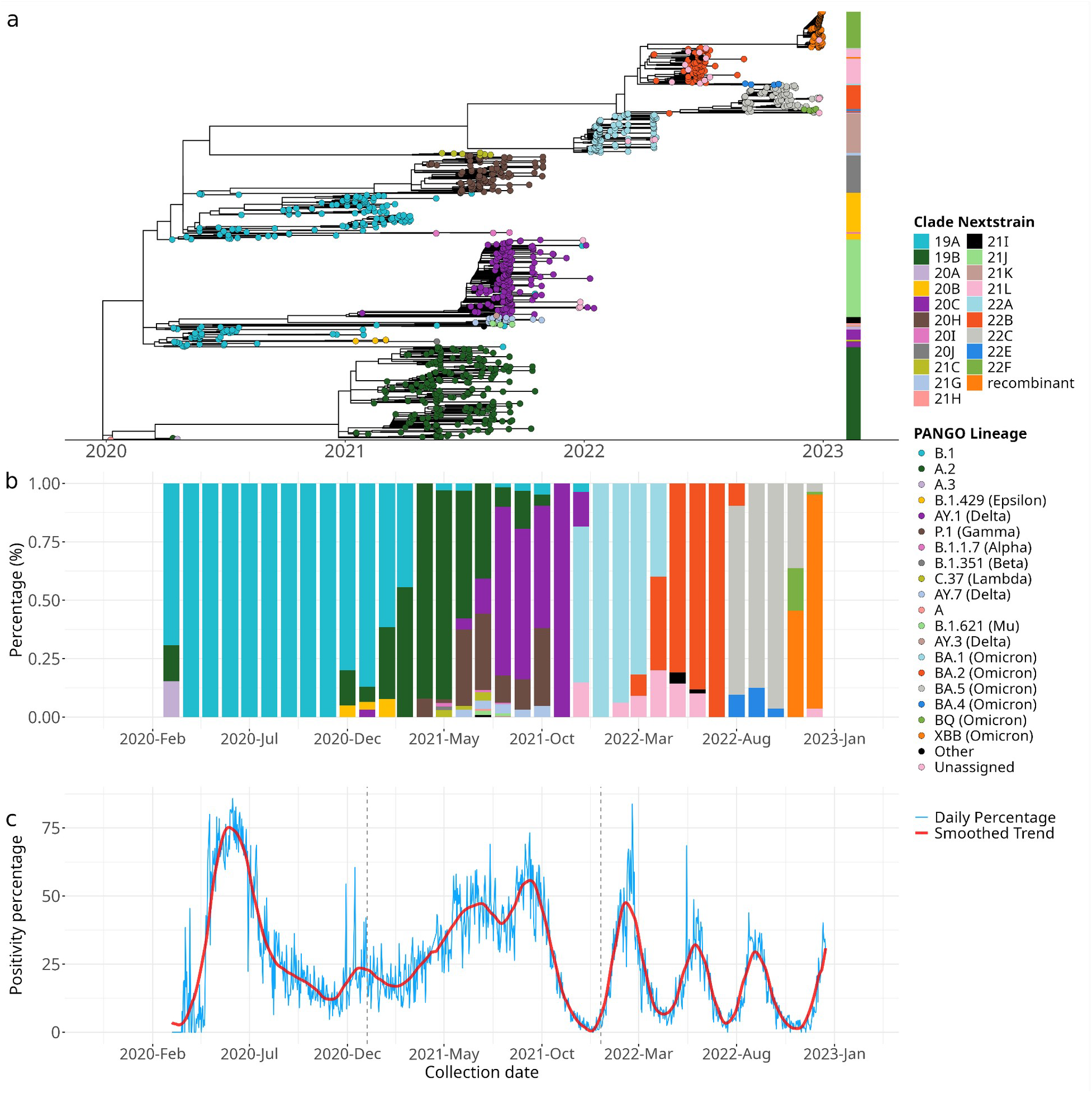
Diversity and temporal dynamics of SARS-CoV-2 lineages in Nicaragua. Lineages are color-coded. Legends present their Nextstrain nomenclature, Pangolin nomenclature, and their WHO Greek-letter designation if they have one based on their status as variants of concern. a. Dated maximum likelihood tree of SARS-CoV-2 sequences from Nicaragua, tips colored according to the Pangolin nomenclature. The Nextstrain classification is indicated by the bar to the right of the tree. b. Proportion of different SARS-CoV-2 lineages over time, where colors correspond to Pangolin lineage, and height indicates the relative frequency of that lineage at a given time. c. Daily positivity percentage (blue line) and smoothed trend (red line) showing changes in COVID-19 positivity rates in Nicaragua from March 2020 to December 2022.

To compare the circulation of variants in Nicaragua to those circulating in the Central American region, we accessed publicly available sequences on the Global Initiative on Sharing All Influenza Data platform, GISAID (https://gisaid.org/) from 2020 to 2022 for a total of 22,530 genomes circulating in Guatemala (n=4,255), Belize (n=222), El Salvador (n=863), Costa Rica (n=9,210), Honduras (n=332), and Panama (n=6,584) (Supplementary Fig. 3).

### Phylogenetic Diversity and Epidemiological Insights

SARS-CoV-2 sequences were assigned to clades using both the Nextstrain platform and nomenclature (https://clades.nextstrain.org/) as well as the Pangolin system and nomenclature (https://pangolin.cog-uk.io/). Nextstrain was able to classify all genomes successfully. However, Pangolin failed to classify 60 genomes, most of which belonged to the Omicron lineage.

Phylogenetic analysis characterizes the genetic diversity circulating in Nicaragua over the study period. Circulating lineages included variants of concern (VOC) as classified by the WHO such as Alpha, Beta, Gamma, Delta, and Omicron. As in the global arena, the circulation of various lineages in Nicaragua changed over time. The initial circulation of SARS-CoV-2 early in 2020 included Pangolin lineage A.1, A.2 and the B.1 lineage, although the latter dominated Nicaragua’s epidemiological landscape throughout 2021 and had the longest residency in the country (Fig. 1a and b). After the B.1 lineage ceased to circulate, the A.2 lineage circulated from late 2020 to October 2021, making it the second longest circulating lineage. While the AY.3 (Delta) lineage was identified by the end of 2020, it was not until mid-2021 that significant circulation of this lineage occurred, dominating the second half of 2021. The AY.1 and AY.2 lineages circulated infrequently. The BA.1 lineage, classified by WHO as Omicron, was identified in December 2021 and by January 2022 had completely replaced the previously circulating AY.3 lineage. Despite the rapid expansion of the BA.1 lineage, it did not circulate as long as previous lineages: after three months, the BA.2 lineage was introduced and became the dominant lineage in the country. During the circulation of BA.2, BA.4 was introduced but did not dominate as its predecessors did. In August 2022, the BA.5 lineage was identified and quickly replaced the circulation of the BA.2 and BA.4 lineages, circulating until November when the BQ and XBB lineages were introduced. By December 2022, the XBB lineage became the dominant lineage in the country. These results, shown in Fig. 1a & b, can be summarized as the dominance of a sole lineage for a period of time in the early days of the pandemic, followed by more rapid turnover in the latter days including the co-circulation of multiple lineages for shorter periods until a new lineage is introduced.

In addition to different lineages circulating over time, we observed increases in the positivity rate, or percentage of cases testing positive, coinciding with the introduction of new lineages to the country. The first positivity peak after the initial introduction of SARS-CoV-2 in Nicaragua occurred in June 2020 coinciding with the fixation of the B.1 lineage (Fig. 1c). In November 2020 another spike in positivity rate corresponded with a resurgence of the A.2 lineage. In 2021, positivity rate slowly increased, peaking in July 2021 when the A.2 and P.1 (Gamma) lineages co-dominated the epidemiological scene, and again when the AY.3 lineage predominated in the second half of 2021. In 2022, four peaks in positivity rate were observed, coinciding with the circulation of the BA.1, BA.2, BA.5, and XBB lineages respectively towards the end of the year.

### Spatio-temporal dynamics of SARS-CoV-2

Nationally, SARS-CoV-2 lineages were identified in all departments of the country. In 2020, the B.1 lineage predominated, with some occurrences of the A.1 and A.2 lineages (Fig 2.a). There were no genomes from the Río San Juan department or the South Caribbean Coast autonomous region during this year. In 2021, various lineages circulated throughout the national territory, predominantly the Delta variant lineages in the Pacific region, with AY.3 being prevalent, along with B1 lineages in some areas, A.2, and P.1 (gamma) (Fig. 2b). Similarly, in the Central region and the Autonomous Caribbean regions, the AY.3 lineage of the variant was most frequent. During 2022, all Omicron variant lineages were widely distributed throughout the country, nearly in equal proportions (Fig. 2c). The department of Managua had the highest number of sequences corresponding to more cases over the three years studied, followed by the northern departments. This pattern remained more or less consistent over the three years.

**Figure 2.**
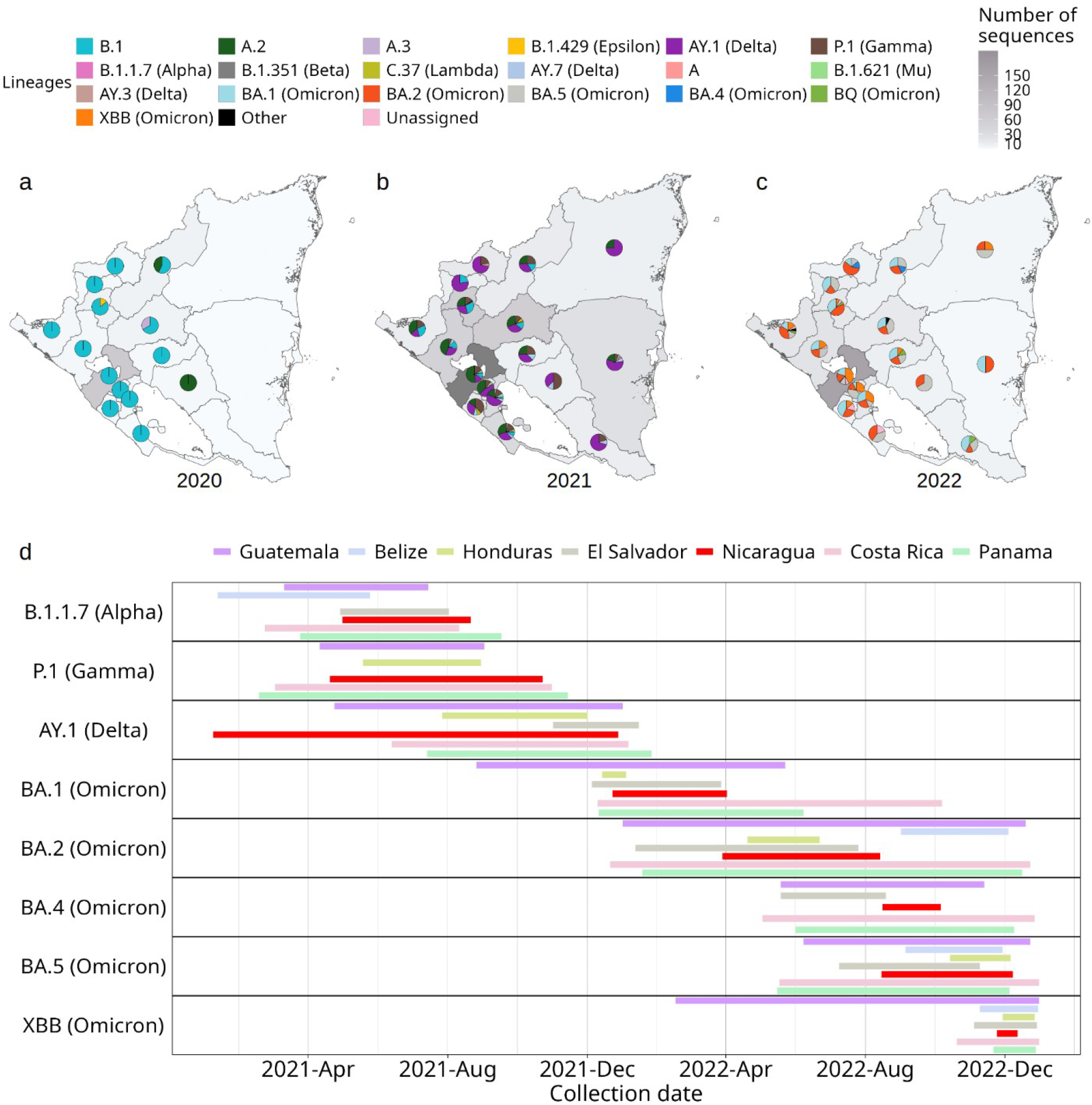
Spatial and temporal distribution of SARS-CoV-2 lineages. Frequency of genomes included in this study and distribution of lineages by department during the years 2020 (a), 2021 (b), and 2022 (c); in all three panels the number of sequences per region is indicated by gray shading. d: Temporal circulation of variants in Central American countries. Lineages are named using Pangolin nomenclature and their WHO Greek-letter designation if they have one.

**Figure 3.**
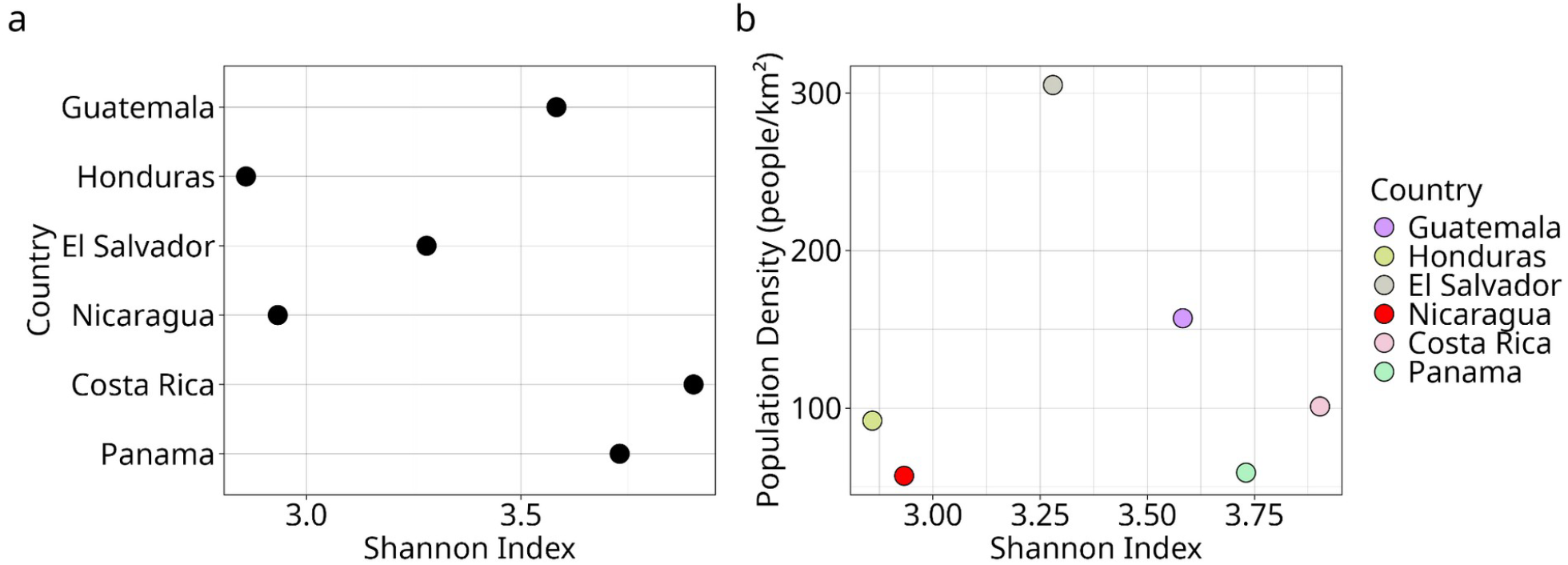
Diversity of Circulating Lineages in Central America: a) Shannon diversity index of SARS-CoV-2 lineages in Central American countries. b) Shannon diversity index versus population density (inhabitants per square kilometer) in Central American countries

An inspection of the earliest sample collection dates of viral genomes in this study in Nicaragua was also conducted (Supplementary table 2). Following the introduction of SARS-COV-2 into the country in March 2020, the earliest viral genomes belong to the B.1, A.1, and A.2 lineages in the departments of Jinotega, Chontales, and Matagalpa, respectively. Similarly, other variants such as B.1.429 (Epsilon), AY.1 (Delta), and P.1 (Gamma) were initially reported in Estelí, RACCN, and Estelí respectively, at different times from December 2020 to April 2021. This pattern of early detection suggests a wide geographical spread of the virus variants across the national territory during the initial phases of the outbreak.

To characterize the circulation of variants in the Central American region by date, 22,530 genomes were downloaded from GISAID. Results indicate that many of the SARS-CoV-2 lineages circulating in the Central American region arrived early on in their global spread (Fig. 2d). Belize, Costa Rica, Guatemala and Panama show the earliest records of B.1.1.7 (Alpha) in the region. Nicaragua, although late to record most of the variants, stands out for the early introduction of the AY.1 (Delta) lineage followed by an extensive period of circulation. Guatemala stands out as the country in the region among the earliest to record several variants, and first to record multiple Omicron variants, including BA.1 and XBB. Several of the variants, and XBB in particular, persisted in Guatemala for longer periods than in other countries, suggesting Guatemala may have been a critical entry point and dispersion hub for strains circulating in Central America. In general, our analysis emphasizes a sequence of introductions not confined to a single country but rather characterized as a mosaic of introductions across various points in Central America.

To further examine how Central American countries have hosted a diversity of variants, we compared them based on their Shannon diversity indices based on the lineages. Costa Rica and Panama hosted the highest diversity of lineages (Fig. 3a). We then plotted the diversity indices of countries against their population density (people per square kilometer) (Fig. 3b). Although no statistically significant association was found between population density and the diversity index, it was observed that Nicaragua and Honduras, which have the lowest population densities, also exhibit the lowest diversity indices. El Salvador and Guatemala, with higher population densities, also display higher diversity of lineages. However, Costa Rica, despite having a population density similar to that of Honduras, shows high diversity. Finally, it was observed that Panama, despite its low population density, has a considerably high diversity of lineages.

Given the extensive range of mutations observed in SARS-CoV-2 genomes, we assessed the top 30 most frequent amino acid substitutions occurring amongst our sequences from Nicaragua by lineage (Fig. 4). Within the A.2 lineage, which circulated extensively throughout much of 2021, a high number of the top 30 amino acid substitutions was identified. This lineage was followed by the Delta variant lineages and B.1, which both had high counts but across fewer of the top 30 substitutions. The Omicron lineages also accumulated high counts for, and notably across the most of the top 30 amino acid substitutions.

**Figure 4.**
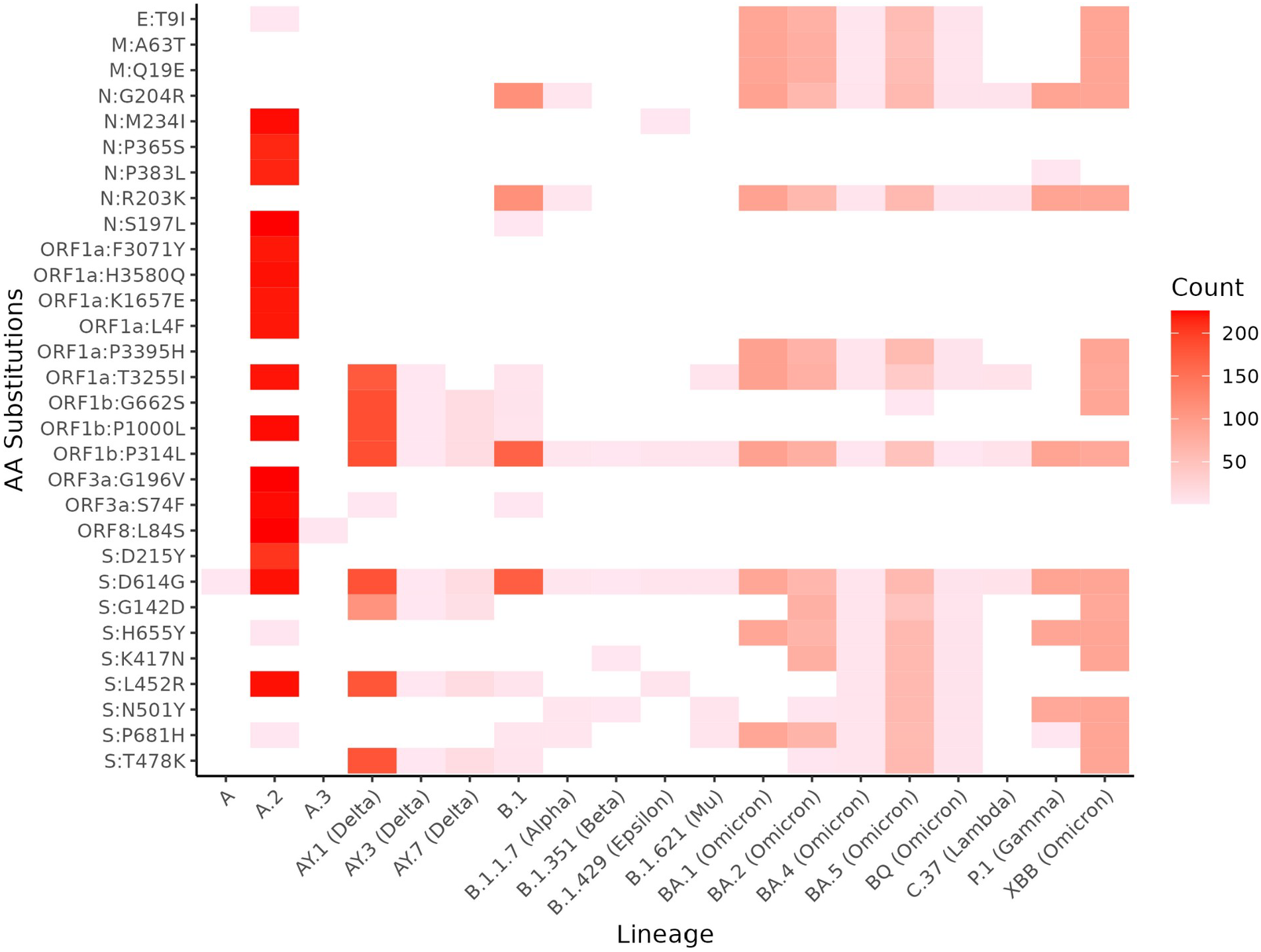
Number of mutations in lineages circulating in Nicaragua. Heatmap indicates the number of times a specific amino acid (aa) substitution (y-axis) was observed amongst our sequences within a given lineage (x-axis). Lineages are named using Pangolin nomenclature and their WHO Greek-letter designation if they have one. Substitution nomenclature follows standard practice - gene: original aa state, gene position, new aa state.

The substitution in the Spike gene, from Aspartic Acid (D) to Glycine (G) at gene position 614 (S:D614G) was present in all genomes except for lineage A.3. Following this, the P314L mutation in the ORF1b gene region was the second most prevalent across the genomes, only absent in lineages A, A.2, and A.3.

We analyzed the association between the top 30 mutations and the likelihood of hospitalization (Fig. 5) and found that there are significant associations between hospitalization and individual mutations in the N region (G204R and R203K), ORF1b region (G662S, P1000L, P314L), and spike protein: D614G, G142D, L452R, and T478K.

**Fig 5.**
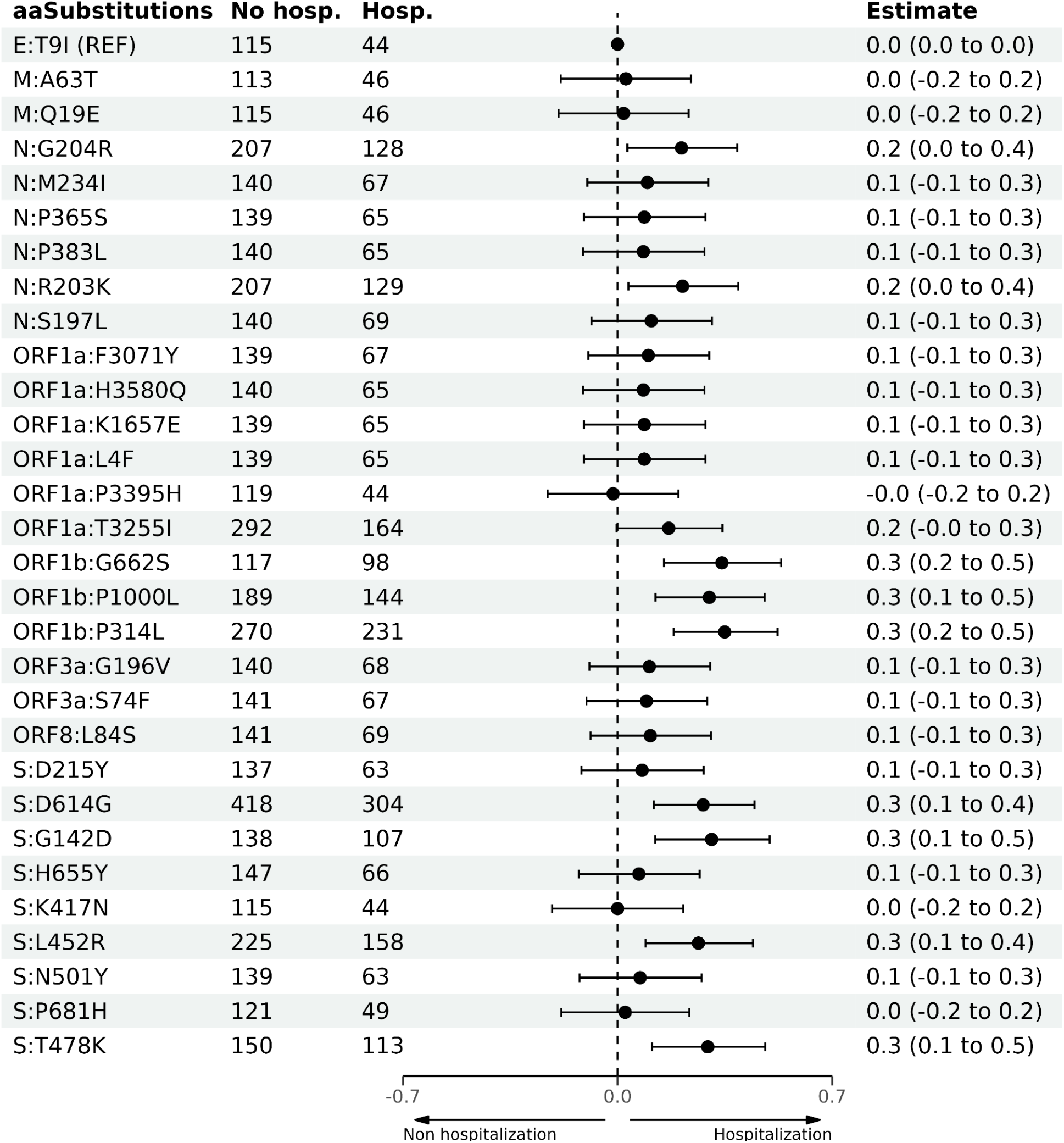
Logistic Regression Model with the 30 Most Frequent Mutations in SARS-CoV-2 Genomes against Hospitalization. The estimate is exponentiated and represents the Odds Ratios (ORs) on a log10 scale where 0 represents no change in risk, values above 0 indicate increased risk, and values below 0 indicate decreased risk of hospitalization

## DISCUSSION

This study represents the first characterization of the genomic diversity of the SARS-CoV-2 lineages circulating in Nicaragua during the first three years since the initial COVID-19 case in March 2020, and is the result of a multi-center collaboration to establish genomic epidemiological surveillance within the country.

During the initial phase of the research, when samples were sent abroad for sequencing, there was no strict stratification by department in the sample selection; however, this was implemented in the second phase when local sequencing was established. We consider it a significant advantage for this study that diagnostic processes for epidemiological surveillance and genomic surveillance are collocated and interconnected. This arrangement has enabled almost immediate access to samples, the capability to perform sequencing in near real-time, and the strategic selection of samples based on the origin of the positive cases.

Similar to the course of the pandemic on a global scale, our data demonstrate patterns of dominance and replacement among different virus lineages over time. During the first year and a half of the pandemic, B.1 and A.2 lineages dominated, even as emerging lineages were being identified in different parts of the world, mainly in European Union countries and American countries like the United States and Brazil^17–19^. The dynamics in Nicaragua during 2021 shifted with the emergence of various lineages and rapid transitions, for example, from Delta (AY.3) to Omicron (BA.1). Throughout the following year, in 2022, various Omicron lineages circulated, leading to changes in positivity patterns.

Genomic surveillance requires viral loads ranging from high to moderate to yield adequate sequence information. Samples reaching the National Virology Laboratory (NVL) in Managua for processing are also at risk of genetic material degradation depending on the travel distance due to transportation time, and may not achieve appropriate cycle threshold (Ct) for sequencing, despite having positive PCR results. The result might be fewer sequences derived from regions farther from Managua. These regions are also typically less populated, resulting in fewer samples sent to the National Center for Disease Control and Prevention (CNDR). Nonetheless, we present a broad geographic representation of sequences across Nicaragua that support our observation of sustained transmission of SARS-CoV-2 involving a nationwide distribution of lineages over the three years of this study.

At a regional level, Nicaragua is part of a multi-country arena of viral exchange involving many of its closest neighbors, including Guatemala, Honduras, and Costa Rica. There is a disparity in the number of sequences available among the countries in the Central American region. Countries such as Belize, El Salvador, and Honduras exhibit a very low number of sequences, which are limited to specific periods. In contrast, Panama and Costa Rica have a significantly higher number of sequences. Nicaragua, while not having as many sequences as its southern neighbors (Costa Rica and Panama), shows a distribution of sequences throughout the duration of this study. Although we failed to demonstrate a link between the population density of a country and the diversity of viral lineages, Certain patterns were observed; countries with low lineage diversity also tend to have low population densities (Honduras and Nicaragua), while countries with high population densities exhibit significant lineage diversity (Guatemala and El Salvador). Others have observed that population dynamics may be a significant modulator of the pandemic’s impact in terms of morbidity and mortality ^20–24^.

According to the data analyzed in this research, the diversity of lineages circulating in Nicaragua is lower compared to the majority of the countries in Central America and the introduction of new lineages occurred later than the majority of the countries in the region. It has been documented that most of the Central American countries, except for Nicaragua, have instituted stringent lockdowns, states of emergency, and states of calamity ^25–30^. This observation suggests that, beyond these measures, there are other potential intrinsic factors in population dynamics that influence the nature of the circulating diversity and establishment of SARS-CoV-2 lineages. We acknowledge that the number of genomic sequences available from various countries depends on their capacity to perform genomic surveillance and sequencing; therefore, this disparity may introduce bias into our results

We identified a relationship between SARS-CoV-2 genomic mutations and hospitalization. The mutations identified in the spike protein, particularly D614G, which have been shown to be associated with hospitalization, are consistent with previous findings suggesting a link between virus transmissibility and disease severity^31–33^. This mutation, due to its prevalence and association with increased infectivity, could indicate a more severe clinical course, leading to a higher number of patients requiring hospitalization. In addition to mutations in spike, our analysis identified the recurrence and association of mutations in the ORF1b region among hospitalized patients, suggesting that alterations in the viral replicative machinery may have a direct impact on virus pathogenesis. The viral polymerase function, encoded by ORF1b, is critical for viral genome replication, and mutations that enhance this process could theoretically lead to increased virulence. Overall, the data are consistent with previous studies showing an association between hospitalization and mutations in specific regions of the SARS-CoV-2 genome^34–36^. However, the observed association between the mutations S:D614G and ORF1b:P314L and hospitalization rates, despite their presence in lineages not typically associated with severe outcomes especially during the last years such as Omicron lineages(pending publication), reflect a complex interplay of viral evolution and host factors. These mutations might be part of an ongoing adaptation of the virus, where emerging lineages are evolving to become less virulent. Additionally, the impact of these mutations on disease severity could be significantly influenced by the development of immunity in the population, either through natural infection or vaccination, which might alter the clinical outcomes of infections with these variants. Furthermore, multifactorial influences including access to healthcare, presence of comorbid conditions, age, and individual immune system responses play critical roles in determining the severity of the disease.

This study was made possible by national viral and clinical surveillance programs in Nicaragua coupled with recent investments in genomic sequencing and analytical capacities. Together, these features contributed new insights into the dynamics of the SARS-CoV-2 pandemic in the country of Nicaragua and the circulation of variants at the national and regional levels.

## METHODS

### Study population and sampling collection

All respiratory swab samples were collected from March 2020 to December 2022 across 15 departments and 2 autonomous regions in Nicaragua, as part of two ongoing studies: the Nicaraguan Pediatric Influenza Cohort Study, and the American-Asian Arbovirus Research and Epidemiological Surveillance (A2CARES) program based in Managua, along with the Nicaraguan national surveillance of respiratory diseases. All the methods were performed in accordance with the relevant guidelines and regulations of the Nicaraguan Ministry of Health.

The Nicaraguan Pediatric Influenza Cohort Study is conducted in the catchment area of the “Sócrates Flores Vivas” Health Center (HCSFV), the principal primary care facility serving neighborhoods along the shores of Lake Xolotlán in Managua, serving a population ranging from 3,693 to 3,947 (Balmaseda et al., 2006; Hammond et al., 2005). The A2CARES program employed a spatial randomization and satellite selection method, devised by Dr. Kathryn Hacker and supervised by Dr. Aubree Gordon at the University of Michigan (Data Management), to randomly select participants from both urban and rural sectors. This program followed 1,000 households (496 rural and 504 urban households), encompassing 1,024 participants in Nejapa (rural) and 1,100 participants in the Camilo Ortega area (urban), with ages ranging from 2 to 80 years old. Additionally, samples from all provinces of the country were collected through the National Surveillance Program of the Nicaraguan Ministry of Health. The National Epidemiological Surveillance System for SARS-CoV-2, implemented by the Ministry of Health in Nicaragua, is a monitoring network whose main objective is to track COVID-19 cases across all regions of the country. This system employs the quantitative Reverse Transcription Polymerase Chain Reaction (qRT-PCR) technique to diagnose SARS-CoV-2 cases, allowing for the tracking of viruses as they emerge. The Ministry of Health is organized into 19 Local Comprehensive Health Care Systems (SILAIS), which have functions of health service provision, administration, and local health governance. The SILAIS are responsible for monitoring COVID-19 cases in their respective localities, managing epidemiological surveillance from primary care centers and local hospitals. In this way, the National SILAIS Network functions as a decentralized structure, where each SILAIS is a fundamental component contributing to the National Epidemiological Surveillance System for SARS-CoV-2, thus enabling the monitoring and surveillance of the SARS-CoV-2 cases at the national level.

All SARS-CoV-2 positive samples identified by qRT-PCR were potential candidates for genomic sequencing. However, due to limited sequencing capabilities within the country during the initial phase of the pandemic, between March 2020 and July 2021, a subset of positive samples was randomly selected and sent to The Icahn School of Medicine at Mount Sinai in New York, USA, for whole-genome sequencing using Illumina technology. The resulting sequences were then sent back to Nicaragua for further analysis. During this period, the selection of the respiratory swabs was not stratified by department; instead, random selection was performed among all positive cases across the country

As a response to the COVID-19 pandemic, genomic surveillance was implemented in Nicaragua starting from July 2021 with the Oxford Nanopore Technology sequencing. As part of this initiative, Respiratory swabs with Ct values ranging from 18 to 30 were randomly selected, stratified by department, based on the number of positive COVID-19 cases reported in the national surveillance data provided by the Ministry of Health. While we aimed to maintain a rigorous stratified sampling process by department, we had no control over the sampling strategies employed by the national health surveillance which could change over time depending of the epidemiological landscape in the country. Additionally, as is typical in sequencing techniques, some samples yielded sequences of poor quality. These were excluded from the analysis to prevent errors in the results. This approach ensured that the selection process was both randomized and reflective of the epidemiological distribution across departments, albeit with noted limitations due to external sampling methodologies and inherent challenges in sequencing quality. From all successfully sequenced samples, those with a coverage of 60% or higher were chosen as a quality parameter.

In total, 566 genomes were obtained from Illumina sequencing in the United States, and 498 genomes were obtained in Nicaragua (336 using the Artic V3 sequencing protocol and 162 using the Oxford Nanopore Technology Midnight protocol).

### RNA extraction and RT-PCR

Samples suspected of SARS-CoV-2 infection were collected during the acute phase of illness and processed at the National Virology Laboratory (NVL). Viral RNA was extracted using the QIAmp viral RNA mini kit (QIAGEN, Germany) following the manufacturer’s instructions. In brief, the process starts with the lysis of viral particles in the sample using a lysis buffer, mixed with a 140µl volume of the sample, to release the viral RNA. The lysate is then combined with a binding buffer, which aids in the attachment of viral RNA to the silica membrane in the spin column. The RNA bound to the membrane is thoroughly washed several times with 500µl of wash buffers to eliminate impurities and other contaminants. The purified RNA is then eluted from the spin column using 60µl of elution buffer, making it suitable for RT-PCR analysis.

Subsequently, qRT-PCR was performed following the standardized Multiplex PCR protocol developed by the Virology Institute at Charité University Hospital (Corman et al., 2020), using the ABI 7500 Fast PCR platform (Applied Biosystems, Foster, CA, USA) for subsequent validation. Throughout the pre-analytical and analytical process, both the samples and viral RNA were handled in type II biosafety cabinets and with all required personal protective equipment.

### Library preparation and Next Generation Sequencing

In Nicaragua, the ONT library was prepared according to the ARTIC V3 or ONT Midnight protocol for PCR tiling, using the rapid barcoding kits SQK-LSK109 or SQK-RBK110.96, respectively. Sequencing was conducted on the MinION MK1B and MinION MK1C platforms.

First, viral RNA was reverse-transcribed using LunaScript RT SuperMix. Subsequently, the genetic material underwent PCR amplification aimed at amplifying almost the entire SARS-CoV-2 genome. For this step, both the ARTIC V3 and Midnight protocols used separate primer pools for overlapping tiled PCR reactions spanning the viral genome. The result was DNA segments ranging from 400 to 1200 base pairs covering the SARS-CoV-2 genome, which were confirmed by 1% agarose gel electrophoresis.

The library preparation process for the ARTIC V3 network protocol includes a DNA end-prep step and subsequent addition of barcodes, after which all individually barcoded samples were pooled into a single library. After a DNA purification step using magnetic beads, this version involves adding crucial molecular adaptors for sequencing. In the ONT Midnight protocol, following amplicon generation, the segments were exposed to tagmentation enzymes that both cut the amplicons and added barcodes. Once the samples were barcoded, a single library was created, and the genetic material was purified using magnetic beads, 70% alcohol and molecular grade water.

AmpureXP purification beads (Beckman Coulter, High Wycombe, UK) were used to clean up the PCR products. The DNA concentration (PCR products and DNA libraries) was measured using the Qubit dsDNA HS Assay Kit for fluorometric DNA measurement (Thermo Fisher Scientific) on a Qubit 4.0 Fluorometer. The final DNA library was then loaded onto a primed MinION flow cell R9.4 (FLO-MIN 106).

### Consensus genomes

During sequencing, the ONT MinKnow software discarded all sequences with a Phred quality score below 8. The resulting fastq files were trimmed and mapped to a reference genome and then assembled to generate consensus sequences using the ARTIC network bioinformatic pipeline (https://artic.network/ncov-2019). For samples sequenced using the ONT Midnight protocol, the bioinformatics pipeline wf-artic was used, incorporated into the ONT EPI2ME platform.

### Phylogenetics analysis

Lineage determination and amino acid substitution analysis were conducted using the Nextstrain platform. Sequence alignments were performed using MAFFT v7.4 and manually inspected with AliView v1.28. The construction of the Maximum Likelihood (ML) phylogenetic tree was carried out using IQTREE version 2.2.2.7 with 1000 bootstrap replicates, employing the ModelFinder feature to select the best nucleotide substitution model. The software TreeTime was used to convert the raw ML tree into a dated tree^37^. It was decided to adopt this approach due to its lower demand on computational resources and its proven satisfactory results in previous studies^38^. The R package ggtree was employed for visualizing and annotating the dated ML tree.

## ETHICS STATEMENT

Protocols for the collection and testing of samples were reviewed and approved by the Institutional Review Boards (IRB) of the University of California, Berkeley (PDCS: 2010-09-2245; PDHS: 2010-06-1649; A2CARES: 2021-03-14191) and the Nicaraguan Ministry of Health (PDCS: CIRE 09/03/07/-008. Ver. 25; PDHS: CIRE 01/10/06-13. Ver. 18; A2CARES: CIRE 02/08/21-114 Ver. 4). Informed consent was obtained from all participants, parents, or legal guardians before enrollment; verbal assent was obtained from children 6-11 y/o and written assent from children ≥12 y/o. All samples from the national surveillance system were de-identified and provided only at the health center level.

## ACKNOWLEDGEMENTS

We would like to express our gratitude to all those who have contributed to this work. We thank the study staff of the PDCSs, A2CARES studies, and the National Virology Laboratory of the Nicaraguan Ministry of Health for their valuable contribution and assistance in carrying out the experiments. We also thank to our colleagues Paúl Cárdenas, Sully Márquez, Belén Prado from the Microbiology Institute of the University San Francisco de Quito, Ecuador who generously volunteered their time and efforts for carrying out the necessary training that generate the knowledge required to carry out the present study.

## AUTHORS CONTRIBUTIONS

JC, EH, AG, SB and AB contributed to the conceptualization of the study. GVA, CC, HM and SA conducted the study. GVA, CC and HM performed the laboratory assays. GVA, HM and SA curated the data. GVA, AG, AB and SB analyzed the data. GVA, AG and JGJ generated the figures and tables. GVA, JGJ and SB wrote the initial manuscript draft, and CC, JC, EH, AG and AB edited the manuscript. All authors reviewed the manuscript for scientific content.

## DATA AVAILABILITY

The genome sequences obtained in this study have been made available through the Global Initiative on Sharing All Influenza Data (GISAID) repository https://gisaid.org/. Accession numbers are provided in Supplementary Table 4

