## Supplementary for "Tracking the genetic diversity of SARS-CoV-2 variants in Nicaragua throughout the COVID-19 Pandemic"

Supplementary figure 1: Distribution of the genomes obtained in Nicaragua grouped by lineage (a), department (b) and sequencing method (c)

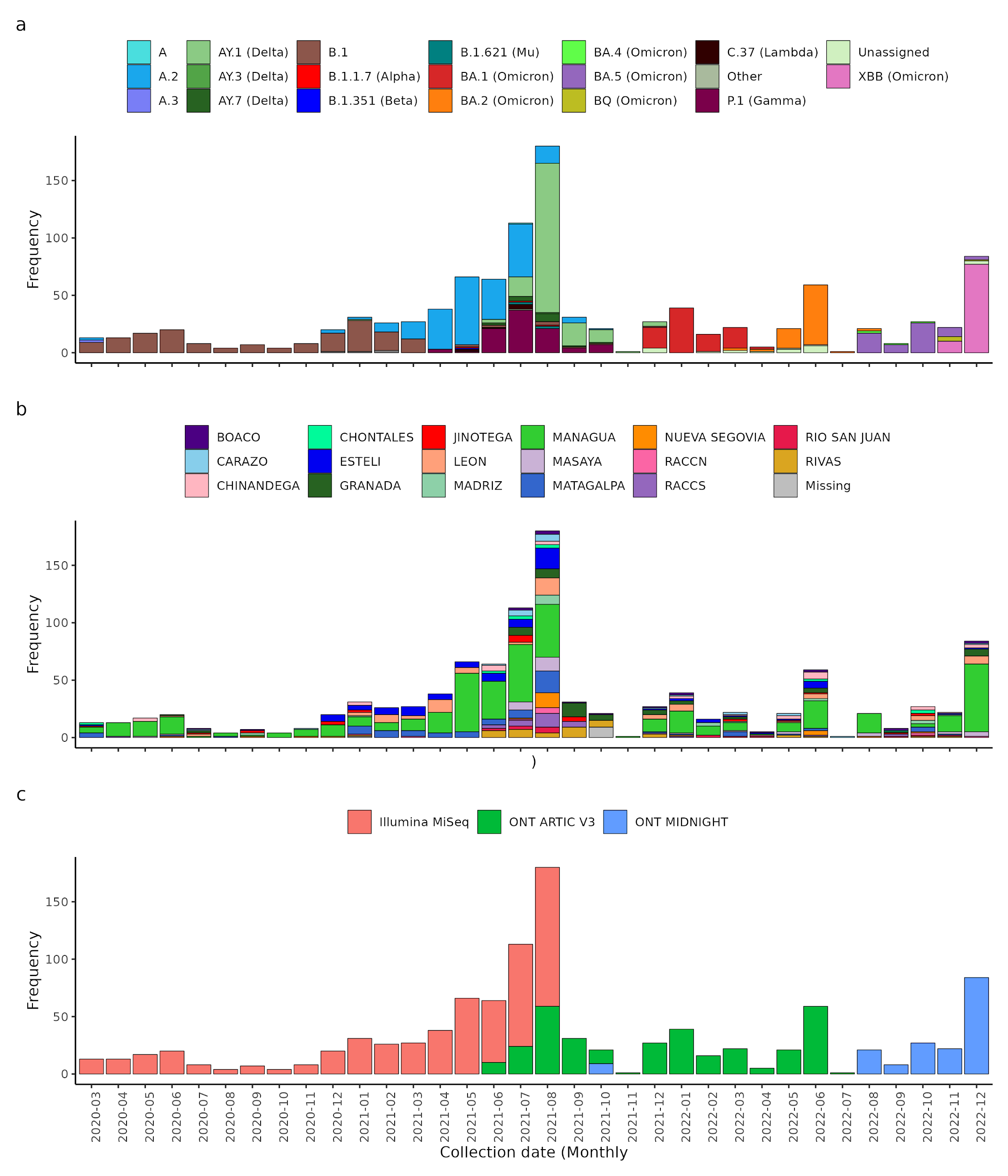

Supplementary figure 2: Distribution of lineages identified in Nicaragua by department.
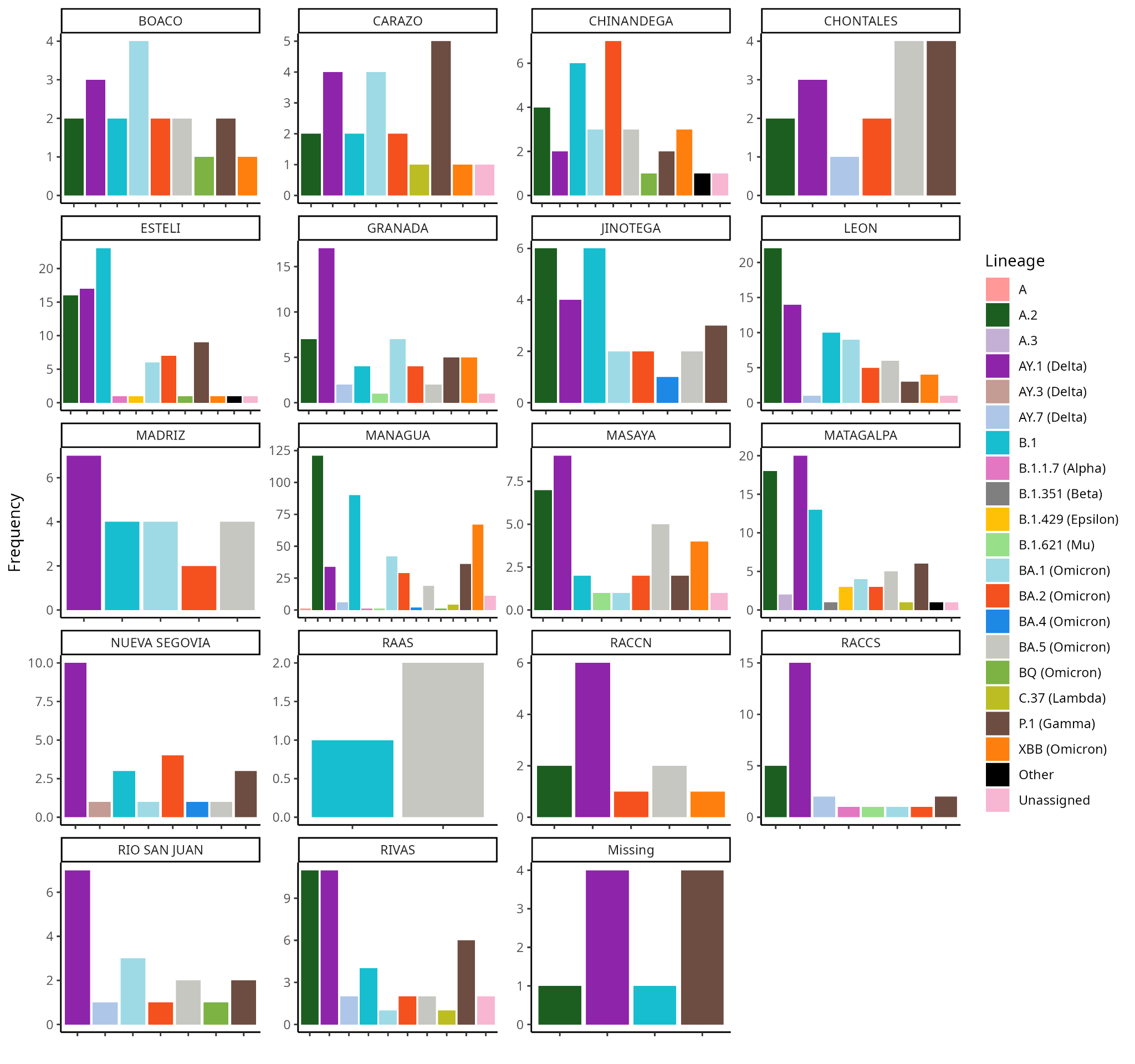

Supplementary figure 3: Frequency of Central American genomes obtained from the GISAID platform distributed per month.

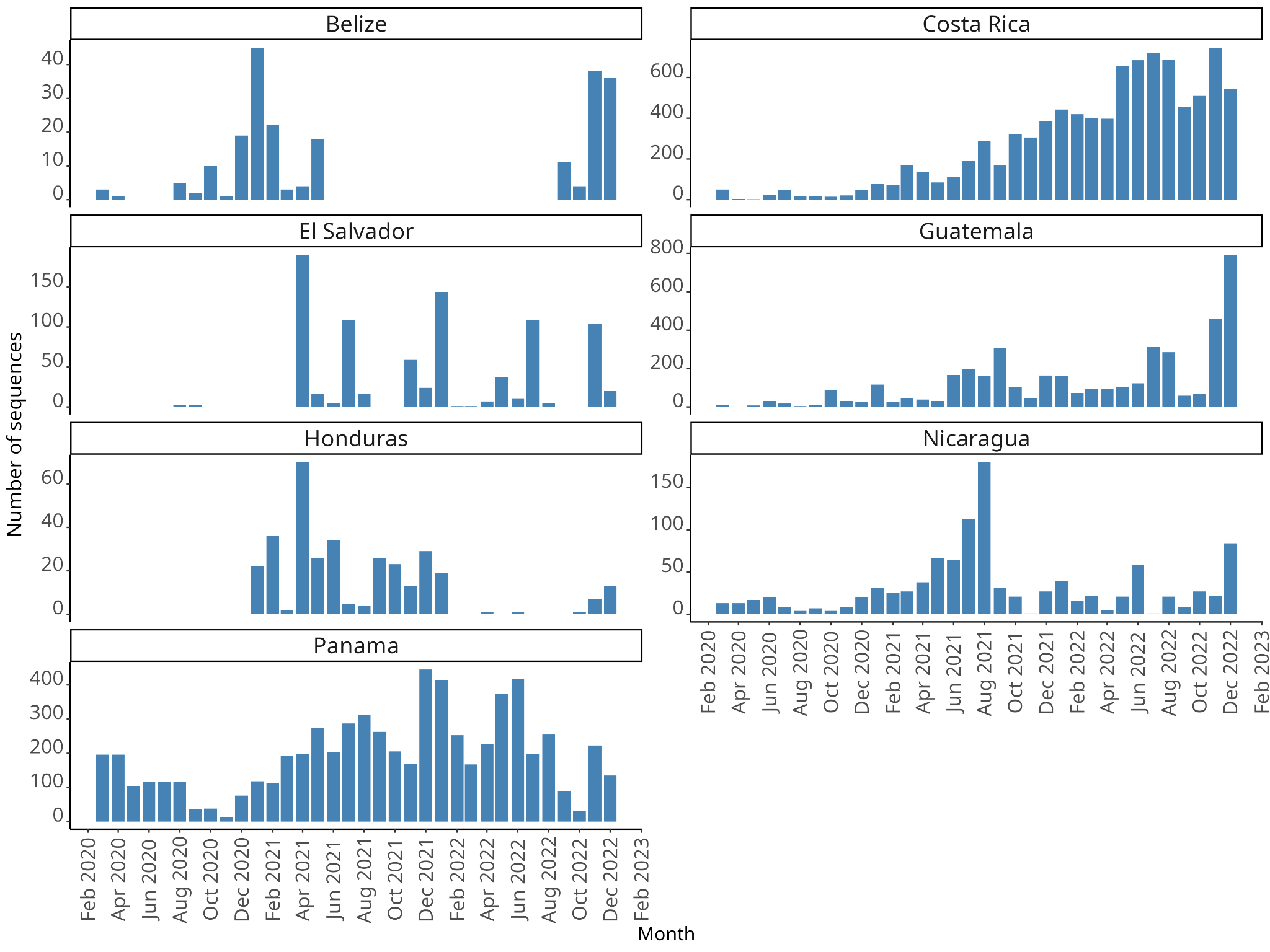

Supplementary Table 1: General summary of the characteristics of the genomes obtained from Nicaragua

| **Lineage** | **Total**  **n(%)** | **Female**  **n(%)** | **Male**  **n(%)** | **Gender Unknown**  **n(%)** | **Hospitalized**  **n(%)** | **Not Hospitalized**  **n(%)** | **Non Hospitalization data**  **n(%)** |
| --- | --- | --- | --- | --- | --- | --- | --- |
| A | 1 (0.09) | 1 (0.09) | 0 (0) | 0 (0) | 0 (0) | 0 (0) | 1 (0.09) |
| A.2 | 226 (21.24) | 131 (12.31) | 95 (8.93) | 0 (0) | 68 (6.39) | 140 (13.16) | 18 (1.69) |
| A.3 | 2 (0.19) | 0 (0) | 2 (0.19) | 0 (0) | 1 (0.09) | 1 (0.09) | 0 (0) |
| AY.1 (Delta) | 187 (17.58) | 111 (10.43) | 76 (7.14) | 0 (0) | 66 (6.2) | 47 (4.42) | 74 (6.95) |
| AY.3 (Delta) | 1 (0.09) | 0 (0) | 1 (0.09) | 0 (0) | 1 (0.09) | 0 (0) | 0 (0) |
| AY.7 (Delta) | 15 (1.41) | 8 (0.75) | 7 (0.66) | 0 (0) | 9 (0.85) | 1 (0.09) | 5 (0.47) |
| B.1 | 171 (16.07) | 90 (8.46) | 81 (7.61) | 0 (0) | 95 (8.93) | 70 (6.58) | 6 (0.56) |
| B.1.1.7 (Alpha) | 3 (0.28) | 1 (0.09) | 2 (0.19) | 0 (0) | 2 (0.19) | 1 (0.09) | 0 (0) |
| B.1.351 (Beta) | 1 (0.09) | 1 (0.09) | 0 (0) | 0 (0) | 0 (0) | 1 (0.09) | 0 (0) |
| B.1.429 (Epsilon) | 4 (0.38) | 1 (0.09) | 3 (0.28) | 0 (0) | 1 (0.09) | 3 (0.28) | 0 (0) |
| B.1.621 (Mu) | 4 (0.38) | 3 (0.28) | 1 (0.09) | 0 (0) | 0 (0) | 4 (0.38) | 0 (0) |
| BA.1 (Omicron) | 92 (8.65) | 46 (4.32) | 44 (4.14) | 2 (0.19) | 1 (0.09) | 7 (0.66) | 84 (7.89) |
| BA.2 (Omicron) | 76 (7.14) | 59 (5.55) | 17 (1.6) | 0 (0) | 5 (0.47) | 10 (0.94) | 61 (5.73) |
| BA.4 (Omicron) | 4 (0.38) | 4 (0.38) | 0 (0) | 0 (0) | 1 (0.09) | 1 (0.09) | 2 (0.19) |
| BA.5 (Omicron) | 61 (5.73) | 42 (3.95) | 19 (1.79) | 0 (0) | 18 (1.69) | 27 (2.54) | 16 (1.5) |
| BQ (Omicron) | 5 (0.47) | 3 (0.28) | 2 (0.19) | 0 (0) | 0 (0) | 5 (0.47) | 0 (0) |
| C.37 (Lambda) | 7 (0.66) | 2 (0.19) | 5 (0.47) | 0 (0) | 0 (0) | 3 (0.28) | 4 (0.38) |
| Other | 3 (0.28) | 3 (0.28) | 0 (0) | 0 (0) | 1 (0.09) | 1 (0.09) | 1 (0.09) |
| P.1 (Gamma) | 94 (8.83) | 50 (4.7) | 44 (4.14) | 0 (0) | 23 (2.16) | 31 (2.91) | 40 (3.76) |
| Unassigned | 20 (1.88) | 7 (0.66) | 13 (1.22) | 0 (0) | 4 (0.38) | 4 (0.38) | 12 (1.13) |
| XBB (Omicron) | 87 (8.18) | 57 (5.36) | 30 (2.82) | 0 (0) | 19 (1.79) | 68 (6.39) | 0 (0) |

Supplementary table 2: Early identification and location of the Nicaraguan genomes.

| **Lineage** | **Early Identification** | **Early Location** |
| --- | --- | --- |
| B.1 | 2020-Mar | JINOTEGA |
| A.2 | 2020-Mar | CHONTALES |
| A.3 | 2020-Mar | MATAGALPA |
| B.1.429 (Epsilon) | 2020-Dec | ESTELÍ |
| AY.1 (Delta) | 2021-Jan | RACCN |
| P.1 (Gamma) | 2021-Apr | ESTELÍ |
| B.1.1.7 (Alpha) | 2021-May | ESTELÍ |
| B.1.351 (Beta) | 2021-May | MATAGALPA |
| C.37 (Lambda) | 2021-May | MANAGUA |
| AY.7 (Delta) | 2021-Jun | RÍO SAN JUAN |
| A | 2021-Jul | MANAGUA |
| B.1.621 (Mu) | 2021-Jul | MASAYA |
| AY.3 (Delta) | 2021-Aug | NUEVA SEGOVIA |
| Unassigned | 2021-Dec | MATAGALPA |
| BA.1 (Omicron) | 2021-Dec | GRANADA |
| BA.2 (Omicron) | 2022-Mar | MANAGUA |
| BA.5 (Omicron) | 2022-Aug | MANAGUA |
| BA.4 (Omicron) | 2022-Aug | MANAGUA |
| BQ (Omicron) | 2022-Nov | RÍO SAN JUAN |
| XBB (Omicron) | 2022-Nov | MANAGUA |

Supplementary table 3: Frequencies and Percentages of Sequences by Department, 2020-2022 in Nicaragua.

| **Department** | **2020** | **2021** | **2022** | **Total** | **Total (%)** |
| --- | --- | --- | --- | --- | --- |
| MANAGUA | 70 | 235 | 160 | 465 | 43,70 |
| ESTELÍ | 7 | 61 | 16 | 84 | 7,89 |
| MATAGALPA | 6 | 59 | 13 | 78 | 7,33 |
| LEÓN | 3 | 50 | 22 | 75 | 7,05 |
| GRANADA | 3 | 36 | 16 | 55 | 5,17 |
| RIVAS | 1 | 36 | 5 | 42 | 3,95 |
| MASAYA | 2 | 19 | 13 | 34 | 3,20 |
| CHINANDEGA | 3 | 11 | 19 | 33 | 3,10 |
| RACCS | 0 | 26 | 2 | 28 | 2,63 |
| JINOTEGA | 7 | 12 | 7 | 26 | 2,44 |
| NUEVA SEGOVIA | 3 | 14 | 7 | 24 | 2,26 |
| CARAZO | 2 | 13 | 7 | 22 | 2,07 |
| MADRIZ | 2 | 9 | 10 | 21 | 1,97 |
| BOACO | 2 | 8 | 9 | 19 | 1,79 |
| RÍO SAN JUAN | 0 | 10 | 7 | 17 | 1,60 |
| CHONTALES | 2 | 8 | 6 | 16 | 1,50 |
| RACCN | 0 | 8 | 4 | 12 | 1,13 |
| RAAS | 1 | 0 | 2 | 3 | 0,28 |
| NO DATA | 0 | 10 | 0 | 10 | 0,94 |

Supplementary table 4: GISAID Accession IDs for 1064 genomes generated in Nicaragua

| Accession ID | Collection date | Country |
| --- | --- | --- |
| EPI_ISL_9159722 | 2020-03-23 | Nicaragua |
| EPI_ISL_9159809 | 2020-03-24 | Nicaragua |
| EPI_ISL_9159790 | 2020-03-24 | Nicaragua |
| EPI_ISL_9159836 | 2020-03-25 | Nicaragua |
| EPI_ISL_9160088 | 2020-03-25 | Nicaragua |
| EPI_ISL_9160084 | 2020-03-25 | Nicaragua |
| EPI_ISL_9159586 | 2020-03-25 | Nicaragua |
| EPI_ISL_9159839 | 2020-03-26 | Nicaragua |
| EPI_ISL_9159840 | 2020-03-27 | Nicaragua |
| EPI_ISL_9159847 | 2020-03-30 | Nicaragua |
| EPI_ISL_9159845 | 2020-03-30 | Nicaragua |
| EPI_ISL_9159848 | 2020-03-31 | Nicaragua |
| EPI_ISL_9159849 | 2020-03-31 | Nicaragua |
| EPI_ISL_9159865 | 2020-04-03 | Nicaragua |
| EPI_ISL_9159879 | 2020-04-08 | Nicaragua |
| EPI_ISL_9159966 | 2020-04-11 | Nicaragua |
| EPI_ISL_9160086 | 2020-04-14 | Nicaragua |
| EPI_ISL_9160083 | 2020-04-14 | Nicaragua |
| EPI_ISL_9160085 | 2020-04-14 | Nicaragua |
| EPI_ISL_9160089 | 2020-04-15 | Nicaragua |
| EPI_ISL_9159768 | 2020-04-21 | Nicaragua |
| EPI_ISL_9159779 | 2020-04-21 | Nicaragua |
| EPI_ISL_9159798 | 2020-04-21 | Nicaragua |
| EPI_ISL_9159843 | 2020-04-27 | Nicaragua |
| EPI_ISL_9159842 | 2020-04-27 | Nicaragua |
| EPI_ISL_9159841 | 2020-04-27 | Nicaragua |
| EPI_ISL_9159862 | 2020-05-04 | Nicaragua |
| EPI_ISL_9159852 | 2020-05-04 | Nicaragua |
| EPI_ISL_9159859 | 2020-05-04 | Nicaragua |
| EPI_ISL_9159856 | 2020-05-04 | Nicaragua |
| EPI_ISL_9159855 | 2020-05-04 | Nicaragua |
| EPI_ISL_10281011 | 2020-05-04 | Nicaragua |
| EPI_ISL_9159853 | 2020-05-04 | Nicaragua |
| EPI_ISL_9159851 | 2020-05-04 | Nicaragua |
| EPI_ISL_9159860 | 2020-05-04 | Nicaragua |
| EPI_ISL_9159858 | 2020-05-04 | Nicaragua |
| EPI_ISL_9159854 | 2020-05-04 | Nicaragua |
| EPI_ISL_9159857 | 2020-05-04 | Nicaragua |
| EPI_ISL_9159861 | 2020-05-04 | Nicaragua |
| EPI_ISL_9159878 | 2020-05-07 | Nicaragua |
| EPI_ISL_9159875 | 2020-05-07 | Nicaragua |
| EPI_ISL_9159876 | 2020-05-07 | Nicaragua |
| EPI_ISL_9159874 | 2020-05-07 | Nicaragua |
| EPI_ISL_9160063 | 2020-06-02 | Nicaragua |
| EPI_ISL_9160066 | 2020-06-02 | Nicaragua |
| EPI_ISL_9160079 | 2020-06-04 | Nicaragua |
| EPI_ISL_9160080 | 2020-06-05 | Nicaragua |
| EPI_ISL_9160081 | 2020-06-05 | Nicaragua |
| EPI_ISL_9159637 | 2020-06-09 | Nicaragua |
| EPI_ISL_9160093 | 2020-06-10 | Nicaragua |
| EPI_ISL_9160092 | 2020-06-10 | Nicaragua |
| EPI_ISL_9160095 | 2020-06-11 | Nicaragua |
| EPI_ISL_9160096 | 2020-06-11 | Nicaragua |
| EPI_ISL_9160103 | 2020-06-15 | Nicaragua |
| EPI_ISL_9159626 | 2020-06-17 | Nicaragua |
| EPI_ISL_9159624 | 2020-06-17 | Nicaragua |
| EPI_ISL_9159636 | 2020-06-19 | Nicaragua |
| EPI_ISL_9159929 | 2020-06-20 | Nicaragua |
| EPI_ISL_9159667 | 2020-06-21 | Nicaragua |
| EPI_ISL_9159669 | 2020-06-23 | Nicaragua |
| EPI_ISL_9159670 | 2020-06-23 | Nicaragua |
| EPI_ISL_9159671 | 2020-06-23 | Nicaragua |
| EPI_ISL_9160002 | 2020-06-29 | Nicaragua |
| EPI_ISL_9159783 | 2020-07-01 | Nicaragua |
| EPI_ISL_9159782 | 2020-07-01 | Nicaragua |
| EPI_ISL_9159834 | 2020-07-03 | Nicaragua |
| EPI_ISL_9159833 | 2020-07-03 | Nicaragua |
| EPI_ISL_9159835 | 2020-07-03 | Nicaragua |
| EPI_ISL_9159837 | 2020-07-07 | Nicaragua |
| EPI_ISL_9159838 | 2020-07-15 | Nicaragua |
| EPI_ISL_9159846 | 2020-07-31 | Nicaragua |
| EPI_ISL_9159590 | 2020-08-02 | Nicaragua |
| EPI_ISL_9159850 | 2020-08-17 | Nicaragua |
| EPI_ISL_9159863 | 2020-08-27 | Nicaragua |
| EPI_ISL_9159864 | 2020-08-27 | Nicaragua |
| EPI_ISL_9160105 | 2020-09-09 | Nicaragua |
| EPI_ISL_9159868 | 2020-09-18 | Nicaragua |
| EPI_ISL_9159867 | 2020-09-18 | Nicaragua |
| EPI_ISL_9159871 | 2020-09-18 | Nicaragua |
| EPI_ISL_9159870 | 2020-09-18 | Nicaragua |
| EPI_ISL_9159869 | 2020-09-18 | Nicaragua |
| EPI_ISL_9159872 | 2020-09-29 | Nicaragua |
| EPI_ISL_9159873 | 2020-10-02 | Nicaragua |
| EPI_ISL_9159633 | 2020-10-23 | Nicaragua |
| EPI_ISL_9159927 | 2020-10-23 | Nicaragua |
| EPI_ISL_9159880 | 2020-10-27 | Nicaragua |
| EPI_ISL_9159882 | 2020-11-07 | Nicaragua |
| EPI_ISL_9159881 | 2020-11-07 | Nicaragua |
| EPI_ISL_9159721 | 2020-11-23 | Nicaragua |
| EPI_ISL_9159883 | 2020-11-24 | Nicaragua |
| EPI_ISL_9159884 | 2020-11-29 | Nicaragua |
| EPI_ISL_9159886 | 2020-11-30 | Nicaragua |
| EPI_ISL_9159885 | 2020-11-30 | Nicaragua |
| EPI_ISL_9159887 | 2020-11-30 | Nicaragua |
| EPI_ISL_9159889 | 2020-12-09 | Nicaragua |
| EPI_ISL_9159888 | 2020-12-09 | Nicaragua |
| EPI_ISL_9159890 | 2020-12-15 | Nicaragua |
| EPI_ISL_9159892 | 2020-12-16 | Nicaragua |
| EPI_ISL_9159893 | 2020-12-16 | Nicaragua |
| EPI_ISL_9159894 | 2020-12-16 | Nicaragua |
| EPI_ISL_9159891 | 2020-12-16 | Nicaragua |
| EPI_ISL_9159895 | 2020-12-16 | Nicaragua |
| EPI_ISL_9159896 | 2020-12-16 | Nicaragua |
| EPI_ISL_9159898 | 2020-12-28 | Nicaragua |
| EPI_ISL_9159897 | 2020-12-28 | Nicaragua |
| EPI_ISL_9159899 | 2020-12-28 | Nicaragua |
| EPI_ISL_9159903 | 2020-12-29 | Nicaragua |
| EPI_ISL_9159902 | 2020-12-29 | Nicaragua |
| EPI_ISL_9159900 | 2020-12-29 | Nicaragua |
| EPI_ISL_9159901 | 2020-12-29 | Nicaragua |
| EPI_ISL_9159904 | 2020-12-30 | Nicaragua |
| EPI_ISL_9159907 | 2020-12-30 | Nicaragua |
| EPI_ISL_9159905 | 2020-12-30 | Nicaragua |
| EPI_ISL_9159906 | 2020-12-30 | Nicaragua |
| EPI_ISL_9159910 | 2021-01-04 | Nicaragua |
| EPI_ISL_9159909 | 2021-01-04 | Nicaragua |
| EPI_ISL_9159908 | 2021-01-04 | Nicaragua |
| EPI_ISL_9159734 | 2021-01-08 | Nicaragua |
| EPI_ISL_9159913 | 2021-01-09 | Nicaragua |
| EPI_ISL_9159912 | 2021-01-09 | Nicaragua |
| EPI_ISL_9159911 | 2021-01-09 | Nicaragua |
| EPI_ISL_9159914 | 2021-01-09 | Nicaragua |
| EPI_ISL_9159916 | 2021-01-10 | Nicaragua |
| EPI_ISL_9159915 | 2021-01-10 | Nicaragua |
| EPI_ISL_9159917 | 2021-01-11 | Nicaragua |
| EPI_ISL_9159918 | 2021-01-11 | Nicaragua |
| EPI_ISL_9159919 | 2021-01-12 | Nicaragua |
| EPI_ISL_9159921 | 2021-01-12 | Nicaragua |
| EPI_ISL_9159920 | 2021-01-12 | Nicaragua |
| EPI_ISL_9159922 | 2021-01-12 | Nicaragua |
| EPI_ISL_9159923 | 2021-01-15 | Nicaragua |
| EPI_ISL_9159924 | 2021-01-15 | Nicaragua |
| EPI_ISL_9159925 | 2021-01-17 | Nicaragua |
| EPI_ISL_9159926 | 2021-01-17 | Nicaragua |
| EPI_ISL_9159627 | 2021-01-19 | Nicaragua |
| EPI_ISL_9159932 | 2021-01-25 | Nicaragua |
| EPI_ISL_9159933 | 2021-01-25 | Nicaragua |
| EPI_ISL_9159937 | 2021-01-26 | Nicaragua |
| EPI_ISL_9159934 | 2021-01-26 | Nicaragua |
| EPI_ISL_9159936 | 2021-01-27 | Nicaragua |
| EPI_ISL_9159935 | 2021-01-27 | Nicaragua |
| EPI_ISL_9159931 | 2021-01-28 | Nicaragua |
| EPI_ISL_9159938 | 2021-01-29 | Nicaragua |
| EPI_ISL_10281012 | 2021-01-30 | Nicaragua |
| EPI_ISL_9159939 | 2021-01-30 | Nicaragua |
| EPI_ISL_9159969 | 2021-02-01 | Nicaragua |
| EPI_ISL_9159940 | 2021-02-02 | Nicaragua |
| EPI_ISL_9159942 | 2021-02-05 | Nicaragua |
| EPI_ISL_9159941 | 2021-02-07 | Nicaragua |
| EPI_ISL_9159945 | 2021-02-08 | Nicaragua |
| EPI_ISL_9159944 | 2021-02-10 | Nicaragua |
| EPI_ISL_9159943 | 2021-02-10 | Nicaragua |
| EPI_ISL_9159950 | 2021-02-10 | Nicaragua |
| EPI_ISL_9159947 | 2021-02-10 | Nicaragua |
| EPI_ISL_9159951 | 2021-02-10 | Nicaragua |
| EPI_ISL_10281013 | 2021-02-11 | Nicaragua |
| EPI_ISL_9159958 | 2021-02-12 | Nicaragua |
| EPI_ISL_9159952 | 2021-02-13 | Nicaragua |
| EPI_ISL_9159948 | 2021-02-13 | Nicaragua |
| EPI_ISL_9159949 | 2021-02-13 | Nicaragua |
| EPI_ISL_9159957 | 2021-02-13 | Nicaragua |
| EPI_ISL_9159954 | 2021-02-14 | Nicaragua |
| EPI_ISL_9159953 | 2021-02-15 | Nicaragua |
| EPI_ISL_9159956 | 2021-02-15 | Nicaragua |
| EPI_ISL_9159960 | 2021-02-21 | Nicaragua |
| EPI_ISL_10281015 | 2021-02-23 | Nicaragua |
| EPI_ISL_9159962 | 2021-02-23 | Nicaragua |
| EPI_ISL_9159964 | 2021-02-24 | Nicaragua |
| EPI_ISL_9159971 | 2021-02-26 | Nicaragua |
| EPI_ISL_9159963 | 2021-02-27 | Nicaragua |
| EPI_ISL_9159968 | 2021-02-28 | Nicaragua |
| EPI_ISL_9159970 | 2021-03-03 | Nicaragua |
| EPI_ISL_9159974 | 2021-03-06 | Nicaragua |
| EPI_ISL_9159733 | 2021-03-08 | Nicaragua |
| EPI_ISL_9159976 | 2021-03-09 | Nicaragua |
| EPI_ISL_9159973 | 2021-03-10 | Nicaragua |
| EPI_ISL_9159975 | 2021-03-10 | Nicaragua |
| EPI_ISL_9159977 | 2021-03-12 | Nicaragua |
| EPI_ISL_9159981 | 2021-03-13 | Nicaragua |
| EPI_ISL_9159980 | 2021-03-15 | Nicaragua |
| EPI_ISL_10281018 | 2021-03-15 | Nicaragua |
| EPI_ISL_9159978 | 2021-03-15 | Nicaragua |
| EPI_ISL_9159984 | 2021-03-17 | Nicaragua |
| EPI_ISL_9159979 | 2021-03-18 | Nicaragua |
| EPI_ISL_10281017 | 2021-03-19 | Nicaragua |
| EPI_ISL_10281016 | 2021-03-19 | Nicaragua |
| EPI_ISL_9159983 | 2021-03-20 | Nicaragua |
| EPI_ISL_9159988 | 2021-03-20 | Nicaragua |
| EPI_ISL_9159985 | 2021-03-21 | Nicaragua |
| EPI_ISL_9159989 | 2021-03-22 | Nicaragua |
| EPI_ISL_9160017 | 2021-03-23 | Nicaragua |
| EPI_ISL_9159982 | 2021-03-23 | Nicaragua |
| EPI_ISL_9159991 | 2021-03-24 | Nicaragua |
| EPI_ISL_10281019 | 2021-03-24 | Nicaragua |
| EPI_ISL_10281031 | 2021-03-26 | Nicaragua |
| EPI_ISL_9159992 | 2021-03-27 | Nicaragua |
| EPI_ISL_9159993 | 2021-03-27 | Nicaragua |
| EPI_ISL_9159994 | 2021-03-29 | Nicaragua |
| EPI_ISL_10281020 | 2021-04-01 | Nicaragua |
| EPI_ISL_9160001 | 2021-04-05 | Nicaragua |
| EPI_ISL_10281021 | 2021-04-08 | Nicaragua |
| EPI_ISL_9160087 | 2021-04-09 | Nicaragua |
| EPI_ISL_9159990 | 2021-04-10 | Nicaragua |
| EPI_ISL_9159999 | 2021-04-10 | Nicaragua |
| EPI_ISL_9160000 | 2021-04-10 | Nicaragua |
| EPI_ISL_9159998 | 2021-04-10 | Nicaragua |
| EPI_ISL_9160101 | 2021-04-11 | Nicaragua |
| EPI_ISL_9160004 | 2021-04-14 | Nicaragua |
| EPI_ISL_10281024 | 2021-04-15 | Nicaragua |
| EPI_ISL_9160003 | 2021-04-16 | Nicaragua |
| EPI_ISL_9160008 | 2021-04-16 | Nicaragua |
| EPI_ISL_10281022 | 2021-04-17 | Nicaragua |
| EPI_ISL_9160009 | 2021-04-17 | Nicaragua |
| EPI_ISL_9160007 | 2021-04-19 | Nicaragua |
| EPI_ISL_10281023 | 2021-04-19 | Nicaragua |
| EPI_ISL_9160010 | 2021-04-19 | Nicaragua |
| EPI_ISL_9160006 | 2021-04-19 | Nicaragua |
| EPI_ISL_9160005 | 2021-04-20 | Nicaragua |
| EPI_ISL_9160031 | 2021-04-21 | Nicaragua |
| EPI_ISL_10281025 | 2021-04-21 | Nicaragua |
| EPI_ISL_9159593 | 2021-04-22 | Nicaragua |
| EPI_ISL_10281028 | 2021-04-22 | Nicaragua |
| EPI_ISL_9160032 | 2021-04-23 | Nicaragua |
| EPI_ISL_9160013 | 2021-04-24 | Nicaragua |
| EPI_ISL_9160012 | 2021-04-24 | Nicaragua |
| EPI_ISL_9160030 | 2021-04-24 | Nicaragua |
| EPI_ISL_9160011 | 2021-04-24 | Nicaragua |
| EPI_ISL_10281026 | 2021-04-24 | Nicaragua |
| EPI_ISL_9160082 | 2021-04-25 | Nicaragua |
| EPI_ISL_10281027 | 2021-04-25 | Nicaragua |
| EPI_ISL_10280991 | 2021-04-27 | Nicaragua |
| EPI_ISL_9160033 | 2021-04-27 | Nicaragua |
| EPI_ISL_10281014 | 2021-04-27 | Nicaragua |
| EPI_ISL_9160025 | 2021-04-29 | Nicaragua |
| EPI_ISL_9160052 | 2021-04-30 | Nicaragua |
| EPI_ISL_9160042 | 2021-04-30 | Nicaragua |
| EPI_ISL_9160021 | 2021-05-01 | Nicaragua |
| EPI_ISL_9160054 | 2021-05-01 | Nicaragua |
| EPI_ISL_9160022 | 2021-05-01 | Nicaragua |
| EPI_ISL_9160029 | 2021-05-01 | Nicaragua |
| EPI_ISL_9160026 | 2021-05-01 | Nicaragua |
| EPI_ISL_9160019 | 2021-05-01 | Nicaragua |
| EPI_ISL_9160023 | 2021-05-01 | Nicaragua |
| EPI_ISL_9160018 | 2021-05-01 | Nicaragua |
| EPI_ISL_9160024 | 2021-05-01 | Nicaragua |
| EPI_ISL_9160027 | 2021-05-01 | Nicaragua |
| EPI_ISL_9160020 | 2021-05-01 | Nicaragua |
| EPI_ISL_9160028 | 2021-05-01 | Nicaragua |
| EPI_ISL_9160102 | 2021-05-02 | Nicaragua |
| EPI_ISL_9160098 | 2021-05-02 | Nicaragua |
| EPI_ISL_9160041 | 2021-05-02 | Nicaragua |
| EPI_ISL_9160034 | 2021-05-02 | Nicaragua |
| EPI_ISL_9160039 | 2021-05-02 | Nicaragua |
| EPI_ISL_9160040 | 2021-05-02 | Nicaragua |
| EPI_ISL_10281029 | 2021-05-02 | Nicaragua |
| EPI_ISL_9160035 | 2021-05-02 | Nicaragua |
| EPI_ISL_9160043 | 2021-05-02 | Nicaragua |
| EPI_ISL_9160036 | 2021-05-02 | Nicaragua |
| EPI_ISL_9160053 | 2021-05-02 | Nicaragua |
| EPI_ISL_9160037 | 2021-05-02 | Nicaragua |
| EPI_ISL_9159972 | 2021-05-03 | Nicaragua |
| EPI_ISL_9160049 | 2021-05-03 | Nicaragua |
| EPI_ISL_9160046 | 2021-05-05 | Nicaragua |
| EPI_ISL_9160045 | 2021-05-05 | Nicaragua |
| EPI_ISL_9160047 | 2021-05-05 | Nicaragua |
| EPI_ISL_9160048 | 2021-05-05 | Nicaragua |
| EPI_ISL_9160044 | 2021-05-05 | Nicaragua |
| EPI_ISL_9160051 | 2021-05-06 | Nicaragua |
| EPI_ISL_9160050 | 2021-05-06 | Nicaragua |
| EPI_ISL_10281030 | 2021-05-06 | Nicaragua |
| EPI_ISL_9160056 | 2021-05-07 | Nicaragua |
| EPI_ISL_9160055 | 2021-05-07 | Nicaragua |
| EPI_ISL_9160062 | 2021-05-07 | Nicaragua |
| EPI_ISL_9160059 | 2021-05-07 | Nicaragua |
| EPI_ISL_9160060 | 2021-05-07 | Nicaragua |
| EPI_ISL_9160057 | 2021-05-07 | Nicaragua |
| EPI_ISL_9160058 | 2021-05-07 | Nicaragua |
| EPI_ISL_9160061 | 2021-05-07 | Nicaragua |
| EPI_ISL_9160070 | 2021-05-08 | Nicaragua |
| EPI_ISL_9160072 | 2021-05-08 | Nicaragua |
| EPI_ISL_9160074 | 2021-05-08 | Nicaragua |
| EPI_ISL_9160073 | 2021-05-08 | Nicaragua |
| EPI_ISL_9160071 | 2021-05-08 | Nicaragua |
| EPI_ISL_9160065 | 2021-05-08 | Nicaragua |
| EPI_ISL_9160068 | 2021-05-08 | Nicaragua |
| EPI_ISL_9160067 | 2021-05-08 | Nicaragua |
| EPI_ISL_9160069 | 2021-05-08 | Nicaragua |
| EPI_ISL_9160064 | 2021-05-08 | Nicaragua |
| EPI_ISL_9159612 | 2021-05-10 | Nicaragua |
| EPI_ISL_9159986 | 2021-05-14 | Nicaragua |
| EPI_ISL_9160016 | 2021-05-18 | Nicaragua |
| EPI_ISL_9159697 | 2021-05-20 | Nicaragua |
| EPI_ISL_9159959 | 2021-05-20 | Nicaragua |
| EPI_ISL_9159653 | 2021-05-22 | Nicaragua |
| EPI_ISL_9159630 | 2021-05-23 | Nicaragua |
| EPI_ISL_9159654 | 2021-05-24 | Nicaragua |
| EPI_ISL_9159634 | 2021-05-26 | Nicaragua |
| EPI_ISL_9159660 | 2021-05-26 | Nicaragua |
| EPI_ISL_9160014 | 2021-05-27 | Nicaragua |
| EPI_ISL_9159997 | 2021-05-28 | Nicaragua |
| EPI_ISL_9160076 | 2021-05-28 | Nicaragua |
| EPI_ISL_9159987 | 2021-05-30 | Nicaragua |
| EPI_ISL_9159635 | 2021-06-01 | Nicaragua |
| EPI_ISL_9159946 | 2021-06-03 | Nicaragua |
| EPI_ISL_17065458 | 2021-06-07 | Nicaragua |
| EPI_ISL_17065460 | 2021-06-07 | Nicaragua |
| EPI_ISL_17065459 | 2021-06-07 | Nicaragua |
| EPI_ISL_9160015 | 2021-06-07 | Nicaragua |
| EPI_ISL_17065461 | 2021-06-08 | Nicaragua |
| EPI_ISL_17065463 | 2021-06-09 | Nicaragua |
| EPI_ISL_17065462 | 2021-06-09 | Nicaragua |
| EPI_ISL_9159632 | 2021-06-09 | Nicaragua |
| EPI_ISL_17065739 | 2021-06-10 | Nicaragua |
| EPI_ISL_17065464 | 2021-06-10 | Nicaragua |
| EPI_ISL_17065465 | 2021-06-10 | Nicaragua |
| EPI_ISL_9160097 | 2021-06-10 | Nicaragua |
| EPI_ISL_9160078 | 2021-06-11 | Nicaragua |
| EPI_ISL_9159866 | 2021-06-11 | Nicaragua |
| EPI_ISL_9159655 | 2021-06-11 | Nicaragua |
| EPI_ISL_9159621 | 2021-06-12 | Nicaragua |
| EPI_ISL_9159601 | 2021-06-12 | Nicaragua |
| EPI_ISL_9159614 | 2021-06-14 | Nicaragua |
| EPI_ISL_9159659 | 2021-06-14 | Nicaragua |
| EPI_ISL_9159623 | 2021-06-16 | Nicaragua |
| EPI_ISL_9159598 | 2021-06-16 | Nicaragua |
| EPI_ISL_9159607 | 2021-06-17 | Nicaragua |
| EPI_ISL_10280986 | 2021-06-17 | Nicaragua |
| EPI_ISL_9159595 | 2021-06-18 | Nicaragua |
| EPI_ISL_9159605 | 2021-06-18 | Nicaragua |
| EPI_ISL_9159603 | 2021-06-18 | Nicaragua |
| EPI_ISL_9159604 | 2021-06-19 | Nicaragua |
| EPI_ISL_9159617 | 2021-06-19 | Nicaragua |
| EPI_ISL_9159615 | 2021-06-20 | Nicaragua |
| EPI_ISL_10280987 | 2021-06-20 | Nicaragua |
| EPI_ISL_9159619 | 2021-06-20 | Nicaragua |
| EPI_ISL_9159610 | 2021-06-20 | Nicaragua |
| EPI_ISL_9159652 | 2021-06-20 | Nicaragua |
| EPI_ISL_9159602 | 2021-06-21 | Nicaragua |
| EPI_ISL_9159622 | 2021-06-21 | Nicaragua |
| EPI_ISL_9159609 | 2021-06-21 | Nicaragua |
| EPI_ISL_9159596 | 2021-06-21 | Nicaragua |
| EPI_ISL_9159597 | 2021-06-21 | Nicaragua |
| EPI_ISL_9159720 | 2021-06-21 | Nicaragua |
| EPI_ISL_9160090 | 2021-06-21 | Nicaragua |
| EPI_ISL_9159620 | 2021-06-22 | Nicaragua |
| EPI_ISL_9159600 | 2021-06-22 | Nicaragua |
| EPI_ISL_9159611 | 2021-06-22 | Nicaragua |
| EPI_ISL_9159594 | 2021-06-22 | Nicaragua |
| EPI_ISL_9160075 | 2021-06-22 | Nicaragua |
| EPI_ISL_9159618 | 2021-06-23 | Nicaragua |
| EPI_ISL_9159606 | 2021-06-23 | Nicaragua |
| EPI_ISL_9159608 | 2021-06-23 | Nicaragua |
| EPI_ISL_9159995 | 2021-06-24 | Nicaragua |
| EPI_ISL_9159599 | 2021-06-24 | Nicaragua |
| EPI_ISL_10280992 | 2021-06-24 | Nicaragua |
| EPI_ISL_9159629 | 2021-06-24 | Nicaragua |
| EPI_ISL_10280985 | 2021-06-25 | Nicaragua |
| EPI_ISL_10281032 | 2021-06-25 | Nicaragua |
| EPI_ISL_9160091 | 2021-06-25 | Nicaragua |
| EPI_ISL_15326871 | 2021-07-12 | Nicaragua |
| EPI_ISL_9159967 | 2021-06-26 | Nicaragua |
| EPI_ISL_9159616 | 2021-06-26 | Nicaragua |
| EPI_ISL_9159642 | 2021-06-28 | Nicaragua |
| EPI_ISL_9159646 | 2021-06-29 | Nicaragua |
| EPI_ISL_9159640 | 2021-06-30 | Nicaragua |
| EPI_ISL_9159648 | 2021-06-30 | Nicaragua |
| EPI_ISL_9160099 | 2021-07-02 | Nicaragua |
| EPI_ISL_9159588 | 2021-07-02 | Nicaragua |
| EPI_ISL_9159645 | 2021-07-04 | Nicaragua |
| EPI_ISL_10280990 | 2021-07-04 | Nicaragua |
| EPI_ISL_9159996 | 2021-07-04 | Nicaragua |
| EPI_ISL_9159649 | 2021-07-05 | Nicaragua |
| EPI_ISL_9159638 | 2021-07-05 | Nicaragua |
| EPI_ISL_9159651 | 2021-07-05 | Nicaragua |
| EPI_ISL_9159668 | 2021-07-05 | Nicaragua |
| EPI_ISL_9160094 | 2021-07-05 | Nicaragua |
| EPI_ISL_17065737 | 2021-07-06 | Nicaragua |
| EPI_ISL_17065446 | 2021-07-06 | Nicaragua |
| EPI_ISL_17065444 | 2021-07-06 | Nicaragua |
| EPI_ISL_17065442 | 2021-07-06 | Nicaragua |
| EPI_ISL_17065447 | 2021-07-06 | Nicaragua |
| EPI_ISL_17065443 | 2021-07-06 | Nicaragua |
| EPI_ISL_17065738 | 2021-07-06 | Nicaragua |
| EPI_ISL_9159647 | 2021-07-06 | Nicaragua |
| EPI_ISL_9159644 | 2021-07-06 | Nicaragua |
| EPI_ISL_9159641 | 2021-07-06 | Nicaragua |
| EPI_ISL_9159650 | 2021-07-06 | Nicaragua |
| EPI_ISL_9160106 | 2021-07-06 | Nicaragua |
| EPI_ISL_17065445 | 2021-07-06 | Nicaragua |
| EPI_ISL_17065453 | 2021-07-07 | Nicaragua |
| EPI_ISL_17065450 | 2021-07-07 | Nicaragua |
| EPI_ISL_17065449 | 2021-07-07 | Nicaragua |
| EPI_ISL_17065451 | 2021-07-07 | Nicaragua |
| EPI_ISL_17065457 | 2021-07-07 | Nicaragua |
| EPI_ISL_17065454 | 2021-07-07 | Nicaragua |
| EPI_ISL_17065456 | 2021-07-07 | Nicaragua |
| EPI_ISL_17065448 | 2021-07-07 | Nicaragua |
| EPI_ISL_17065452 | 2021-07-07 | Nicaragua |
| EPI_ISL_17065455 | 2021-07-07 | Nicaragua |
| EPI_ISL_9159639 | 2021-07-07 | Nicaragua |
| EPI_ISL_9159666 | 2021-07-07 | Nicaragua |
| EPI_ISL_9159767 | 2021-07-08 | Nicaragua |
| EPI_ISL_9159731 | 2021-07-08 | Nicaragua |
| EPI_ISL_9159643 | 2021-07-08 | Nicaragua |
| EPI_ISL_9159589 | 2021-07-08 | Nicaragua |
| EPI_ISL_9160100 | 2021-07-08 | Nicaragua |
| EPI_ISL_9159591 | 2021-07-10 | Nicaragua |
| EPI_ISL_9159592 | 2021-07-11 | Nicaragua |
| EPI_ISL_17065466 | 2021-07-12 | Nicaragua |
| EPI_ISL_9159613 | 2021-07-12 | Nicaragua |
| EPI_ISL_9159930 | 2021-07-12 | Nicaragua |
| EPI_ISL_9159965 | 2021-07-12 | Nicaragua |
| EPI_ISL_17065740 | 2021-07-13 | Nicaragua |
| EPI_ISL_9159657 | 2021-07-13 | Nicaragua |
| EPI_ISL_9159656 | 2021-07-13 | Nicaragua |
| EPI_ISL_17065467 | 2021-07-13 | Nicaragua |
| EPI_ISL_9159587 | 2021-07-13 | Nicaragua |
| EPI_ISL_17065469 | 2021-07-14 | Nicaragua |
| EPI_ISL_17065468 | 2021-07-14 | Nicaragua |
| EPI_ISL_9159661 | 2021-07-14 | Nicaragua |
| EPI_ISL_17065471 | 2021-07-15 | Nicaragua |
| EPI_ISL_17065470 | 2021-07-15 | Nicaragua |
| EPI_ISL_9159665 | 2021-07-15 | Nicaragua |
| EPI_ISL_9159662 | 2021-07-16 | Nicaragua |
| EPI_ISL_9159928 | 2021-07-16 | Nicaragua |
| EPI_ISL_9159663 | 2021-07-16 | Nicaragua |
| EPI_ISL_9159688 | 2021-07-17 | Nicaragua |
| EPI_ISL_9159683 | 2021-07-18 | Nicaragua |
| EPI_ISL_9159684 | 2021-07-19 | Nicaragua |
| EPI_ISL_9159679 | 2021-07-19 | Nicaragua |
| EPI_ISL_9159664 | 2021-07-19 | Nicaragua |
| EPI_ISL_10280993 | 2021-07-19 | Nicaragua |
| EPI_ISL_10280994 | 2021-07-19 | Nicaragua |
| EPI_ISL_9159682 | 2021-07-20 | Nicaragua |
| EPI_ISL_9159690 | 2021-07-21 | Nicaragua |
| EPI_ISL_9159681 | 2021-07-21 | Nicaragua |
| EPI_ISL_9159700 | 2021-07-21 | Nicaragua |
| EPI_ISL_10280995 | 2021-07-22 | Nicaragua |
| EPI_ISL_9159676 | 2021-07-23 | Nicaragua |
| EPI_ISL_9159698 | 2021-07-23 | Nicaragua |
| EPI_ISL_9159685 | 2021-07-23 | Nicaragua |
| EPI_ISL_9160104 | 2021-07-23 | Nicaragua |
| EPI_ISL_9159585 | 2021-07-23 | Nicaragua |
| EPI_ISL_9159678 | 2021-07-24 | Nicaragua |
| EPI_ISL_9159673 | 2021-07-24 | Nicaragua |
| EPI_ISL_9159675 | 2021-07-24 | Nicaragua |
| EPI_ISL_9159704 | 2021-07-24 | Nicaragua |
| EPI_ISL_9159686 | 2021-07-25 | Nicaragua |
| EPI_ISL_9159699 | 2021-07-25 | Nicaragua |
| EPI_ISL_9159713 | 2021-07-25 | Nicaragua |
| EPI_ISL_9159687 | 2021-07-26 | Nicaragua |
| EPI_ISL_9159705 | 2021-07-26 | Nicaragua |
| EPI_ISL_10280996 | 2021-07-26 | Nicaragua |
| EPI_ISL_9159693 | 2021-07-26 | Nicaragua |
| EPI_ISL_9159707 | 2021-07-26 | Nicaragua |
| EPI_ISL_9159701 | 2021-07-26 | Nicaragua |
| EPI_ISL_9159680 | 2021-07-27 | Nicaragua |
| EPI_ISL_9159677 | 2021-07-27 | Nicaragua |
| EPI_ISL_9159708 | 2021-07-27 | Nicaragua |
| EPI_ISL_9159714 | 2021-07-27 | Nicaragua |
| EPI_ISL_9159692 | 2021-07-27 | Nicaragua |
| EPI_ISL_9159696 | 2021-07-27 | Nicaragua |
| EPI_ISL_9159695 | 2021-07-27 | Nicaragua |
| EPI_ISL_9160077 | 2021-07-27 | Nicaragua |
| EPI_ISL_9159727 | 2021-07-27 | Nicaragua |
| EPI_ISL_9159702 | 2021-07-27 | Nicaragua |
| EPI_ISL_9159703 | 2021-07-27 | Nicaragua |
| EPI_ISL_9159711 | 2021-07-28 | Nicaragua |
| EPI_ISL_9159689 | 2021-07-28 | Nicaragua |
| EPI_ISL_9159706 | 2021-07-28 | Nicaragua |
| EPI_ISL_10280997 | 2021-07-28 | Nicaragua |
| EPI_ISL_9159710 | 2021-07-28 | Nicaragua |
| EPI_ISL_9159694 | 2021-07-28 | Nicaragua |
| EPI_ISL_9159674 | 2021-07-29 | Nicaragua |
| EPI_ISL_9159712 | 2021-07-30 | Nicaragua |
| EPI_ISL_9159691 | 2021-07-30 | Nicaragua |
| EPI_ISL_9159730 | 2021-07-31 | Nicaragua |
| EPI_ISL_9159844 | 2021-07-31 | Nicaragua |
| EPI_ISL_9159747 | 2021-08-01 | Nicaragua |
| EPI_ISL_9159737 | 2021-08-01 | Nicaragua |
| EPI_ISL_9159715 | 2021-08-01 | Nicaragua |
| EPI_ISL_9159719 | 2021-08-01 | Nicaragua |
| EPI_ISL_9159732 | 2021-08-01 | Nicaragua |
| EPI_ISL_9159718 | 2021-08-01 | Nicaragua |
| EPI_ISL_9159716 | 2021-08-01 | Nicaragua |
| EPI_ISL_9159717 | 2021-08-01 | Nicaragua |
| EPI_ISL_9159799 | 2021-08-02 | Nicaragua |
| EPI_ISL_9159729 | 2021-08-02 | Nicaragua |
| EPI_ISL_9159726 | 2021-08-02 | Nicaragua |
| EPI_ISL_9159753 | 2021-08-03 | Nicaragua |
| EPI_ISL_9159631 | 2021-08-03 | Nicaragua |
| EPI_ISL_9159758 | 2021-08-03 | Nicaragua |
| EPI_ISL_10280998 | 2021-08-03 | Nicaragua |
| EPI_ISL_9159724 | 2021-08-03 | Nicaragua |
| EPI_ISL_9159781 | 2021-08-04 | Nicaragua |
| EPI_ISL_9159723 | 2021-08-04 | Nicaragua |
| EPI_ISL_9159728 | 2021-08-04 | Nicaragua |
| EPI_ISL_9159628 | 2021-08-04 | Nicaragua |
| EPI_ISL_9159736 | 2021-08-05 | Nicaragua |
| EPI_ISL_9159778 | 2021-08-05 | Nicaragua |
| EPI_ISL_10281000 | 2021-08-05 | Nicaragua |
| EPI_ISL_10280999 | 2021-08-05 | Nicaragua |
| EPI_ISL_9159725 | 2021-08-05 | Nicaragua |
| EPI_ISL_9159780 | 2021-08-06 | Nicaragua |
| EPI_ISL_9159735 | 2021-08-06 | Nicaragua |
| EPI_ISL_9159748 | 2021-08-07 | Nicaragua |
| EPI_ISL_10280989 | 2021-08-07 | Nicaragua |
| EPI_ISL_9159955 | 2021-08-07 | Nicaragua |
| EPI_ISL_9159755 | 2021-08-08 | Nicaragua |
| EPI_ISL_9159749 | 2021-08-08 | Nicaragua |
| EPI_ISL_9159750 | 2021-08-08 | Nicaragua |
| EPI_ISL_17065505 | 2021-08-10 | Nicaragua |
| EPI_ISL_9159757 | 2021-08-10 | Nicaragua |
| EPI_ISL_9159745 | 2021-08-10 | Nicaragua |
| EPI_ISL_9159746 | 2021-08-10 | Nicaragua |
| EPI_ISL_17065511 | 2021-08-11 | Nicaragua |
| EPI_ISL_17065748 | 2021-08-11 | Nicaragua |
| EPI_ISL_17065510 | 2021-08-11 | Nicaragua |
| EPI_ISL_17065746 | 2021-08-11 | Nicaragua |
| EPI_ISL_17065506 | 2021-08-11 | Nicaragua |
| EPI_ISL_17065509 | 2021-08-11 | Nicaragua |
| EPI_ISL_17065507 | 2021-08-11 | Nicaragua |
| EPI_ISL_17065508 | 2021-08-11 | Nicaragua |
| EPI_ISL_17065745 | 2021-08-11 | Nicaragua |
| EPI_ISL_17065747 | 2021-08-11 | Nicaragua |
| EPI_ISL_10281002 | 2021-08-11 | Nicaragua |
| EPI_ISL_9159658 | 2021-08-11 | Nicaragua |
| EPI_ISL_9159739 | 2021-08-11 | Nicaragua |
| EPI_ISL_9159741 | 2021-08-11 | Nicaragua |
| EPI_ISL_9159742 | 2021-08-11 | Nicaragua |
| EPI_ISL_9159740 | 2021-08-11 | Nicaragua |
| EPI_ISL_9159743 | 2021-08-11 | Nicaragua |
| EPI_ISL_9159744 | 2021-08-11 | Nicaragua |
| EPI_ISL_10280988 | 2021-08-11 | Nicaragua |
| EPI_ISL_9159738 | 2021-08-11 | Nicaragua |
| EPI_ISL_9159751 | 2021-08-12 | Nicaragua |
| EPI_ISL_9159756 | 2021-08-12 | Nicaragua |
| EPI_ISL_9159764 | 2021-08-12 | Nicaragua |
| EPI_ISL_9159765 | 2021-08-12 | Nicaragua |
| EPI_ISL_9159672 | 2021-08-12 | Nicaragua |
| EPI_ISL_17065513 | 2021-08-13 | Nicaragua |
| EPI_ISL_17065512 | 2021-08-13 | Nicaragua |
| EPI_ISL_17065752 | 2021-08-13 | Nicaragua |
| EPI_ISL_17065750 | 2021-08-13 | Nicaragua |
| EPI_ISL_17065751 | 2021-08-13 | Nicaragua |
| EPI_ISL_17065749 | 2021-08-13 | Nicaragua |
| EPI_ISL_10281003 | 2021-08-13 | Nicaragua |
| EPI_ISL_9159754 | 2021-08-13 | Nicaragua |
| EPI_ISL_10281001 | 2021-08-13 | Nicaragua |
| EPI_ISL_9159752 | 2021-08-13 | Nicaragua |
| EPI_ISL_9159766 | 2021-08-13 | Nicaragua |
| EPI_ISL_9159762 | 2021-08-13 | Nicaragua |
| EPI_ISL_17065515 | 2021-08-14 | Nicaragua |
| EPI_ISL_17065514 | 2021-08-14 | Nicaragua |
| EPI_ISL_9159763 | 2021-08-14 | Nicaragua |
| EPI_ISL_9159759 | 2021-08-14 | Nicaragua |
| EPI_ISL_17065472 | 2021-08-15 | Nicaragua |
| EPI_ISL_17065517 | 2021-08-15 | Nicaragua |
| EPI_ISL_17065753 | 2021-08-15 | Nicaragua |
| EPI_ISL_9159828 | 2021-08-15 | Nicaragua |
| EPI_ISL_17065516 | 2021-08-15 | Nicaragua |
| EPI_ISL_9159760 | 2021-08-15 | Nicaragua |
| EPI_ISL_17065480 | 2021-08-16 | Nicaragua |
| EPI_ISL_17065478 | 2021-08-16 | Nicaragua |
| EPI_ISL_17065475 | 2021-08-16 | Nicaragua |
| EPI_ISL_17065473 | 2021-08-16 | Nicaragua |
| EPI_ISL_17065479 | 2021-08-16 | Nicaragua |
| EPI_ISL_17065482 | 2021-08-16 | Nicaragua |
| EPI_ISL_17065481 | 2021-08-16 | Nicaragua |
| EPI_ISL_17065741 | 2021-08-16 | Nicaragua |
| EPI_ISL_17065474 | 2021-08-16 | Nicaragua |
| EPI_ISL_17065484 | 2021-08-16 | Nicaragua |
| EPI_ISL_17065483 | 2021-08-16 | Nicaragua |
| EPI_ISL_17065477 | 2021-08-16 | Nicaragua |
| EPI_ISL_17065476 | 2021-08-16 | Nicaragua |
| EPI_ISL_9159774 | 2021-08-16 | Nicaragua |
| EPI_ISL_9159775 | 2021-08-16 | Nicaragua |
| EPI_ISL_9159769 | 2021-08-16 | Nicaragua |
| EPI_ISL_9159771 | 2021-08-16 | Nicaragua |
| EPI_ISL_9159761 | 2021-08-16 | Nicaragua |
| EPI_ISL_9159776 | 2021-08-16 | Nicaragua |
| EPI_ISL_9159772 | 2021-08-17 | Nicaragua |
| EPI_ISL_9159777 | 2021-08-17 | Nicaragua |
| EPI_ISL_17065486 | 2021-08-19 | Nicaragua |
| EPI_ISL_17065485 | 2021-08-19 | Nicaragua |
| EPI_ISL_9159830 | 2021-08-19 | Nicaragua |
| EPI_ISL_9159832 | 2021-08-19 | Nicaragua |
| EPI_ISL_9159831 | 2021-08-19 | Nicaragua |
| EPI_ISL_17065491 | 2021-08-20 | Nicaragua |
| EPI_ISL_17065487 | 2021-08-20 | Nicaragua |
| EPI_ISL_17065500 | 2021-08-20 | Nicaragua |
| EPI_ISL_17065499 | 2021-08-20 | Nicaragua |
| EPI_ISL_17065743 | 2021-08-20 | Nicaragua |
| EPI_ISL_17065495 | 2021-08-20 | Nicaragua |
| EPI_ISL_17065501 | 2021-08-20 | Nicaragua |
| EPI_ISL_17065492 | 2021-08-20 | Nicaragua |
| EPI_ISL_17065498 | 2021-08-20 | Nicaragua |
| EPI_ISL_17065493 | 2021-08-20 | Nicaragua |
| EPI_ISL_17065502 | 2021-08-20 | Nicaragua |
| EPI_ISL_17065490 | 2021-08-21 | Nicaragua |
| EPI_ISL_17065497 | 2021-08-20 | Nicaragua |
| EPI_ISL_17065494 | 2021-08-20 | Nicaragua |
| EPI_ISL_17065496 | 2021-08-20 | Nicaragua |
| EPI_ISL_17065742 | 2021-08-20 | Nicaragua |
| EPI_ISL_17065488 | 2021-08-20 | Nicaragua |
| EPI_ISL_17065489 | 2021-08-20 | Nicaragua |
| EPI_ISL_9159813 | 2021-08-21 | Nicaragua |
| EPI_ISL_9159804 | 2021-08-21 | Nicaragua |
| EPI_ISL_9159819 | 2021-08-21 | Nicaragua |
| EPI_ISL_9159820 | 2021-08-21 | Nicaragua |
| EPI_ISL_9159812 | 2021-08-21 | Nicaragua |
| EPI_ISL_9159823 | 2021-08-21 | Nicaragua |
| EPI_ISL_9159811 | 2021-08-21 | Nicaragua |
| EPI_ISL_10281007 | 2021-08-21 | Nicaragua |
| EPI_ISL_9159807 | 2021-08-21 | Nicaragua |
| EPI_ISL_9159810 | 2021-08-21 | Nicaragua |
| EPI_ISL_9159808 | 2021-08-21 | Nicaragua |
| EPI_ISL_10281009 | 2021-08-21 | Nicaragua |
| EPI_ISL_9159822 | 2021-08-21 | Nicaragua |
| EPI_ISL_9159797 | 2021-08-21 | Nicaragua |
| EPI_ISL_9159795 | 2021-08-21 | Nicaragua |
| EPI_ISL_9159806 | 2021-08-21 | Nicaragua |
| EPI_ISL_9159814 | 2021-08-21 | Nicaragua |
| EPI_ISL_9159801 | 2021-08-21 | Nicaragua |
| EPI_ISL_9159793 | 2021-08-21 | Nicaragua |
| EPI_ISL_9159821 | 2021-08-21 | Nicaragua |
| EPI_ISL_17065744 | 2021-08-22 | Nicaragua |
| EPI_ISL_17065504 | 2021-08-22 | Nicaragua |
| EPI_ISL_17065503 | 2021-08-22 | Nicaragua |
| EPI_ISL_9159829 | 2021-08-22 | Nicaragua |
| EPI_ISL_9159827 | 2021-08-22 | Nicaragua |
| EPI_ISL_9159791 | 2021-08-22 | Nicaragua |
| EPI_ISL_9159816 | 2021-08-22 | Nicaragua |
| EPI_ISL_10281004 | 2021-08-22 | Nicaragua |
| EPI_ISL_9159794 | 2021-08-22 | Nicaragua |
| EPI_ISL_9159802 | 2021-08-22 | Nicaragua |
| EPI_ISL_9159818 | 2021-08-22 | Nicaragua |
| EPI_ISL_9159824 | 2021-08-22 | Nicaragua |
| EPI_ISL_9159817 | 2021-08-22 | Nicaragua |
| EPI_ISL_9159800 | 2021-08-22 | Nicaragua |
| EPI_ISL_10281008 | 2021-08-22 | Nicaragua |
| EPI_ISL_9159826 | 2021-08-22 | Nicaragua |
| EPI_ISL_9159803 | 2021-08-22 | Nicaragua |
| EPI_ISL_9159825 | 2021-08-22 | Nicaragua |
| EPI_ISL_9159815 | 2021-08-22 | Nicaragua |
| EPI_ISL_10281006 | 2021-08-22 | Nicaragua |
| EPI_ISL_9159792 | 2021-08-22 | Nicaragua |
| EPI_ISL_9159805 | 2021-08-22 | Nicaragua |
| EPI_ISL_9159773 | 2021-08-23 | Nicaragua |
| EPI_ISL_9159770 | 2021-08-23 | Nicaragua |
| EPI_ISL_10281010 | 2021-08-25 | Nicaragua |
| EPI_ISL_9159785 | 2021-08-25 | Nicaragua |
| EPI_ISL_9159784 | 2021-08-25 | Nicaragua |
| EPI_ISL_9159787 | 2021-08-25 | Nicaragua |
| EPI_ISL_9159788 | 2021-08-25 | Nicaragua |
| EPI_ISL_9159789 | 2021-08-25 | Nicaragua |
| EPI_ISL_10281005 | 2021-08-25 | Nicaragua |
| EPI_ISL_9159786 | 2021-08-25 | Nicaragua |
| EPI_ISL_17065754 | 2021-09-06 | Nicaragua |
| EPI_ISL_17065524 | 2021-09-06 | Nicaragua |
| EPI_ISL_17065523 | 2021-09-06 | Nicaragua |
| EPI_ISL_17065525 | 2021-09-07 | Nicaragua |
| EPI_ISL_17065522 | 2021-09-20 | Nicaragua |
| EPI_ISL_17065518 | 2021-09-23 | Nicaragua |
| EPI_ISL_17065521 | 2021-09-23 | Nicaragua |
| EPI_ISL_17065519 | 2021-09-23 | Nicaragua |
| EPI_ISL_17065520 | 2021-09-23 | Nicaragua |
| EPI_ISL_17065535 | 2021-09-25 | Nicaragua |
| EPI_ISL_17065529 | 2021-09-25 | Nicaragua |
| EPI_ISL_17065530 | 2021-09-25 | Nicaragua |
| EPI_ISL_17065527 | 2021-09-25 | Nicaragua |
| EPI_ISL_17065528 | 2021-09-25 | Nicaragua |
| EPI_ISL_17065534 | 2021-09-25 | Nicaragua |
| EPI_ISL_17065526 | 2021-09-25 | Nicaragua |
| EPI_ISL_17065536 | 2021-09-26 | Nicaragua |
| EPI_ISL_17065757 | 2021-09-26 | Nicaragua |
| EPI_ISL_17065538 | 2021-09-26 | Nicaragua |
| EPI_ISL_17065537 | 2021-09-26 | Nicaragua |
| EPI_ISL_17065756 | 2021-09-26 | Nicaragua |
| EPI_ISL_17065531 | 2021-09-28 | Nicaragua |
| EPI_ISL_17065532 | 2021-09-28 | Nicaragua |
| EPI_ISL_17065755 | 2021-09-28 | Nicaragua |
| EPI_ISL_17065533 | 2021-09-28 | Nicaragua |
| EPI_ISL_17065541 | 2021-09-29 | Nicaragua |
| EPI_ISL_17065758 | 2021-09-29 | Nicaragua |
| EPI_ISL_17065540 | 2021-09-29 | Nicaragua |
| EPI_ISL_17065539 | 2021-09-29 | Nicaragua |
| EPI_ISL_17065760 | 2021-09-30 | Nicaragua |
| EPI_ISL_17065759 | 2021-09-30 | Nicaragua |
| EPI_ISL_17065427 | 2021-12-05 | Nicaragua |
| EPI_ISL_17065428 | 2021-12-07 | Nicaragua |
| EPI_ISL_17065734 | 2021-12-07 | Nicaragua |
| EPI_ISL_17065429 | 2021-12-10 | Nicaragua |
| EPI_ISL_17065430 | 2021-12-11 | Nicaragua |
| EPI_ISL_17065722 | 2021-12-16 | Nicaragua |
| EPI_ISL_17065345 | 2021-12-18 | Nicaragua |
| EPI_ISL_17065346 | 2021-12-23 | Nicaragua |
| EPI_ISL_17065347 | 2021-12-23 | Nicaragua |
| EPI_ISL_17065348 | 2021-12-25 | Nicaragua |
| EPI_ISL_14011384 | 2021-12-22 | Nicaragua |
| EPI_ISL_17065349 | 2021-12-28 | Nicaragua |
| EPI_ISL_17065336 | 2021-12-29 | Nicaragua |
| EPI_ISL_17065335 | 2021-12-29 | Nicaragua |
| EPI_ISL_17065350 | 2021-12-29 | Nicaragua |
| EPI_ISL_17065352 | 2021-12-30 | Nicaragua |
| EPI_ISL_17065351 | 2021-12-30 | Nicaragua |
| EPI_ISL_17065353 | 2021-12-31 | Nicaragua |
| EPI_ISL_17065354 | 2021-12-31 | Nicaragua |
| EPI_ISL_17065339 | 2021-12-31 | Nicaragua |
| EPI_ISL_17065340 | 2021-12-31 | Nicaragua |
| EPI_ISL_17065337 | 2021-12-31 | Nicaragua |
| EPI_ISL_17065342 | 2022-01-02 | Nicaragua |
| EPI_ISL_17065338 | 2021-12-31 | Nicaragua |
| EPI_ISL_17065341 | 2021-12-31 | Nicaragua |
| EPI_ISL_15326732 | 2021-12-31 | Nicaragua |
| EPI_ISL_17065718 | 2021-12-31 | Nicaragua |
| EPI_ISL_17065327 | 2022-01-01 | Nicaragua |
| EPI_ISL_17065334 | 2022-01-02 | Nicaragua |
| EPI_ISL_17065343 | 2022-01-02 | Nicaragua |
| EPI_ISL_17065344 | 2022-01-03 | Nicaragua |
| EPI_ISL_17065717 | 2022-01-04 | Nicaragua |
| EPI_ISL_17065328 | 2022-01-04 | Nicaragua |
| EPI_ISL_17065332 | 2022-01-05 | Nicaragua |
| EPI_ISL_17065331 | 2022-01-05 | Nicaragua |
| EPI_ISL_17065333 | 2022-01-05 | Nicaragua |
| EPI_ISL_17065371 | 2022-01-05 | Nicaragua |
| EPI_ISL_17065330 | 2022-01-05 | Nicaragua |
| EPI_ISL_17065329 | 2022-01-05 | Nicaragua |
| EPI_ISL_17065360 | 2022-01-06 | Nicaragua |
| EPI_ISL_17065358 | 2022-01-06 | Nicaragua |
| EPI_ISL_17065359 | 2022-01-06 | Nicaragua |
| EPI_ISL_17065362 | 2022-01-06 | Nicaragua |
| EPI_ISL_17065364 | 2022-01-06 | Nicaragua |
| EPI_ISL_17065361 | 2022-01-06 | Nicaragua |
| EPI_ISL_17065356 | 2022-01-06 | Nicaragua |
| EPI_ISL_17065365 | 2022-01-06 | Nicaragua |
| EPI_ISL_17065355 | 2022-01-06 | Nicaragua |
| EPI_ISL_17065366 | 2022-01-06 | Nicaragua |
| EPI_ISL_17065363 | 2022-01-06 | Nicaragua |
| EPI_ISL_17065357 | 2022-01-06 | Nicaragua |
| EPI_ISL_17065368 | 2022-01-07 | Nicaragua |
| EPI_ISL_17065369 | 2022-01-07 | Nicaragua |
| EPI_ISL_17065723 | 2022-01-07 | Nicaragua |
| EPI_ISL_17065367 | 2022-01-07 | Nicaragua |
| EPI_ISL_17065370 | 2022-01-08 | Nicaragua |
| EPI_ISL_17065853 | 2022-01-18 | Nicaragua |
| EPI_ISL_17065848 | 2022-01-22 | Nicaragua |
| EPI_ISL_17065401 | 2022-01-27 | Nicaragua |
| EPI_ISL_17065405 | 2022-01-29 | Nicaragua |
| EPI_ISL_17065846 | 2022-01-29 | Nicaragua |
| EPI_ISL_17065404 | 2022-01-29 | Nicaragua |
| EPI_ISL_17065852 | 2022-01-30 | Nicaragua |
| EPI_ISL_17065847 | 2022-01-29 | Nicaragua |
| EPI_ISL_17065406 | 2022-02-04 | Nicaragua |
| EPI_ISL_17065407 | 2022-02-06 | Nicaragua |
| EPI_ISL_17065408 | 2022-02-06 | Nicaragua |
| EPI_ISL_17065393 | 2022-02-12 | Nicaragua |
| EPI_ISL_17065392 | 2022-02-12 | Nicaragua |
| EPI_ISL_17065394 | 2022-02-16 | Nicaragua |
| EPI_ISL_17065731 | 2022-02-17 | Nicaragua |
| EPI_ISL_17065399 | 2022-02-18 | Nicaragua |
| EPI_ISL_17065732 | 2022-02-18 | Nicaragua |
| EPI_ISL_17065402 | 2022-02-18 | Nicaragua |
| EPI_ISL_17065403 | 2022-02-18 | Nicaragua |
| EPI_ISL_17065400 | 2022-02-18 | Nicaragua |
| EPI_ISL_17065397 | 2022-02-18 | Nicaragua |
| EPI_ISL_17065396 | 2022-02-18 | Nicaragua |
| EPI_ISL_17065398 | 2022-02-18 | Nicaragua |
| EPI_ISL_17065395 | 2022-02-18 | Nicaragua |
| EPI_ISL_17065770 | 2022-03-28 | Nicaragua |
| EPI_ISL_17065556 | 2022-03-28 | Nicaragua |
| EPI_ISL_17065555 | 2022-03-28 | Nicaragua |
| EPI_ISL_17065566 | 2022-03-28 | Nicaragua |
| EPI_ISL_17065557 | 2022-03-28 | Nicaragua |
| EPI_ISL_17065563 | 2022-03-29 | Nicaragua |
| EPI_ISL_17065772 | 2022-03-29 | Nicaragua |
| EPI_ISL_17065569 | 2022-03-29 | Nicaragua |
| EPI_ISL_17065573 | 2022-03-29 | Nicaragua |
| EPI_ISL_17065567 | 2022-03-29 | Nicaragua |
| EPI_ISL_17065574 | 2022-03-29 | Nicaragua |
| EPI_ISL_17065774 | 2022-03-29 | Nicaragua |
| EPI_ISL_17065572 | 2022-03-29 | Nicaragua |
| EPI_ISL_17065571 | 2022-03-29 | Nicaragua |
| EPI_ISL_17065564 | 2022-03-30 | Nicaragua |
| EPI_ISL_17065570 | 2022-03-30 | Nicaragua |
| EPI_ISL_17065773 | 2022-03-30 | Nicaragua |
| EPI_ISL_17065565 | 2022-03-30 | Nicaragua |
| EPI_ISL_17065568 | 2022-03-30 | Nicaragua |
| EPI_ISL_17065561 | 2022-03-31 | Nicaragua |
| EPI_ISL_17065771 | 2022-03-31 | Nicaragua |
| EPI_ISL_17065562 | 2022-03-31 | Nicaragua |
| EPI_ISL_17065559 | 2022-04-02 | Nicaragua |
| EPI_ISL_17065560 | 2022-04-02 | Nicaragua |
| EPI_ISL_17065558 | 2022-04-02 | Nicaragua |
| EPI_ISL_17065720 | 2022-04-22 | Nicaragua |
| EPI_ISL_17065721 | 2022-04-26 | Nicaragua |
| EPI_ISL_17065441 | 2022-05-14 | Nicaragua |
| EPI_ISL_17065736 | 2022-05-16 | Nicaragua |
| EPI_ISL_17065440 | 2022-05-20 | Nicaragua |
| EPI_ISL_17065439 | 2022-05-21 | Nicaragua |
| EPI_ISL_17065434 | 2022-05-21 | Nicaragua |
| EPI_ISL_17065431 | 2022-05-21 | Nicaragua |
| EPI_ISL_17065438 | 2022-05-21 | Nicaragua |
| EPI_ISL_17065435 | 2022-05-21 | Nicaragua |
| EPI_ISL_17065735 | 2022-05-21 | Nicaragua |
| EPI_ISL_17065433 | 2022-05-21 | Nicaragua |
| EPI_ISL_17065432 | 2022-05-21 | Nicaragua |
| EPI_ISL_17065437 | 2022-05-21 | Nicaragua |
| EPI_ISL_17065436 | 2022-05-21 | Nicaragua |
| EPI_ISL_17065836 | 2022-05-25 | Nicaragua |
| EPI_ISL_17065725 | 2022-05-27 | Nicaragua |
| EPI_ISL_17065724 | 2022-05-27 | Nicaragua |
| EPI_ISL_17065372 | 2022-05-28 | Nicaragua |
| EPI_ISL_17065374 | 2022-05-30 | Nicaragua |
| EPI_ISL_17065375 | 2022-05-30 | Nicaragua |
| EPI_ISL_17065373 | 2022-05-30 | Nicaragua |
| EPI_ISL_17065376 | 2022-05-31 | Nicaragua |
| EPI_ISL_17065377 | 2022-06-01 | Nicaragua |
| EPI_ISL_17065833 | 2022-06-01 | Nicaragua |
| EPI_ISL_17065832 | 2022-06-02 | Nicaragua |
| EPI_ISL_17065726 | 2022-06-02 | Nicaragua |
| EPI_ISL_17065728 | 2022-06-02 | Nicaragua |
| EPI_ISL_17065378 | 2022-06-02 | Nicaragua |
| EPI_ISL_17065727 | 2022-06-02 | Nicaragua |
| EPI_ISL_17065838 | 2022-06-02 | Nicaragua |
| EPI_ISL_17065379 | 2022-06-04 | Nicaragua |
| EPI_ISL_17065381 | 2022-06-04 | Nicaragua |
| EPI_ISL_17065380 | 2022-06-04 | Nicaragua |
| EPI_ISL_17065383 | 2022-06-06 | Nicaragua |
| EPI_ISL_17065382 | 2022-06-06 | Nicaragua |
| EPI_ISL_17065729 | 2022-06-07 | Nicaragua |
| EPI_ISL_17065782 | 2022-06-07 | Nicaragua |
| EPI_ISL_17065385 | 2022-06-07 | Nicaragua |
| EPI_ISL_17065386 | 2022-06-07 | Nicaragua |
| EPI_ISL_17065834 | 2022-06-07 | Nicaragua |
| EPI_ISL_17065594 | 2022-06-07 | Nicaragua |
| EPI_ISL_17065596 | 2022-06-07 | Nicaragua |
| EPI_ISL_17065592 | 2022-06-07 | Nicaragua |
| EPI_ISL_17065384 | 2022-06-07 | Nicaragua |
| EPI_ISL_17065593 | 2022-06-07 | Nicaragua |
| EPI_ISL_17065597 | 2022-06-07 | Nicaragua |
| EPI_ISL_17065591 | 2022-06-07 | Nicaragua |
| EPI_ISL_17065595 | 2022-06-07 | Nicaragua |
| EPI_ISL_17065590 | 2022-06-07 | Nicaragua |
| EPI_ISL_17065389 | 2022-06-08 | Nicaragua |
| EPI_ISL_17065730 | 2022-06-08 | Nicaragua |
| EPI_ISL_17065388 | 2022-06-08 | Nicaragua |
| EPI_ISL_17065837 | 2022-06-08 | Nicaragua |
| EPI_ISL_17065387 | 2022-06-08 | Nicaragua |
| EPI_ISL_17065839 | 2022-06-08 | Nicaragua |
| EPI_ISL_17065835 | 2022-06-08 | Nicaragua |
| EPI_ISL_17065391 | 2022-06-09 | Nicaragua |
| EPI_ISL_17065831 | 2022-06-09 | Nicaragua |
| EPI_ISL_17065390 | 2022-06-09 | Nicaragua |
| EPI_ISL_17065586 | 2022-06-10 | Nicaragua |
| EPI_ISL_17065585 | 2022-06-10 | Nicaragua |
| EPI_ISL_17065589 | 2022-06-10 | Nicaragua |
| EPI_ISL_17065588 | 2022-06-10 | Nicaragua |
| EPI_ISL_17065781 | 2022-06-10 | Nicaragua |
| EPI_ISL_17065587 | 2022-06-10 | Nicaragua |
| EPI_ISL_17065780 | 2022-06-13 | Nicaragua |
| EPI_ISL_17065779 | 2022-06-14 | Nicaragua |
| EPI_ISL_17065778 | 2022-06-14 | Nicaragua |
| EPI_ISL_17065583 | 2022-06-15 | Nicaragua |
| EPI_ISL_17065777 | 2022-06-15 | Nicaragua |
| EPI_ISL_17065584 | 2022-06-15 | Nicaragua |
| EPI_ISL_17065581 | 2022-06-16 | Nicaragua |
| EPI_ISL_17065582 | 2022-06-16 | Nicaragua |
| EPI_ISL_17065580 | 2022-06-17 | Nicaragua |
| EPI_ISL_17065577 | 2022-06-17 | Nicaragua |
| EPI_ISL_17065579 | 2022-06-17 | Nicaragua |
| EPI_ISL_17065578 | 2022-06-17 | Nicaragua |
| EPI_ISL_17065776 | 2022-06-17 | Nicaragua |
| EPI_ISL_17065775 | 2022-06-20 | Nicaragua |
| EPI_ISL_17065576 | 2022-06-23 | Nicaragua |
| EPI_ISL_17065575 | 2022-06-23 | Nicaragua |
| EPI_ISL_13993660 | 2022-07-01 | Nicaragua |
| EPI_ISL_15326994 | 2022-08-04 | Nicaragua |
| EPI_ISL_15326990 | 2022-08-14 | Nicaragua |
| EPI_ISL_17062238 | 2022-09-08 | Nicaragua |
| EPI_ISL_17062239 | 2022-09-10 | Nicaragua |
| EPI_ISL_17063865 | 2022-09-14 | Nicaragua |
| EPI_ISL_17063866 | 2022-09-15 | Nicaragua |
| EPI_ISL_17062240 | 2022-09-18 | Nicaragua |
| EPI_ISL_17062241 | 2022-09-26 | Nicaragua |
| EPI_ISL_17062242 | 2022-09-28 | Nicaragua |
| EPI_ISL_17062243 | 2022-09-29 | Nicaragua |
| EPI_ISL_17062244 | 2022-10-01 | Nicaragua |
| EPI_ISL_17062245 | 2022-10-02 | Nicaragua |
| EPI_ISL_17062246 | 2022-10-03 | Nicaragua |
| EPI_ISL_17063867 | 2022-10-05 | Nicaragua |
| EPI_ISL_17062247 | 2022-10-06 | Nicaragua |
| EPI_ISL_17062248 | 2022-10-06 | Nicaragua |
| EPI_ISL_17062249 | 2022-10-06 | Nicaragua |
| EPI_ISL_17062250 | 2022-10-08 | Nicaragua |
| EPI_ISL_17062251 | 2022-10-08 | Nicaragua |
| EPI_ISL_17062252 | 2022-10-10 | Nicaragua |
| EPI_ISL_17062253 | 2022-10-10 | Nicaragua |
| EPI_ISL_17062254 | 2022-10-11 | Nicaragua |
| EPI_ISL_17062255 | 2022-10-14 | Nicaragua |
| EPI_ISL_17062256 | 2022-10-14 | Nicaragua |
| EPI_ISL_17062257 | 2022-10-14 | Nicaragua |
| EPI_ISL_17062258 | 2022-10-17 | Nicaragua |
| EPI_ISL_17062259 | 2022-10-18 | Nicaragua |
| EPI_ISL_17062260 | 2022-10-18 | Nicaragua |
| EPI_ISL_17062261 | 2022-10-18 | Nicaragua |
| EPI_ISL_17062262 | 2022-10-19 | Nicaragua |
| EPI_ISL_17062263 | 2022-10-20 | Nicaragua |
| EPI_ISL_17062264 | 2022-10-23 | Nicaragua |
| EPI_ISL_17062265 | 2022-10-23 | Nicaragua |
| EPI_ISL_17062266 | 2022-10-25 | Nicaragua |
| EPI_ISL_17062267 | 2022-10-27 | Nicaragua |
| EPI_ISL_17062268 | 2022-10-28 | Nicaragua |
| EPI_ISL_17062269 | 2022-10-30 | Nicaragua |
| EPI_ISL_17062270 | 2022-11-01 | Nicaragua |
| EPI_ISL_17062271 | 2022-11-01 | Nicaragua |
| EPI_ISL_17062272 | 2022-11-04 | Nicaragua |
| EPI_ISL_17062273 | 2022-11-08 | Nicaragua |
| EPI_ISL_17062274 | 2022-11-14 | Nicaragua |
| EPI_ISL_17062275 | 2022-11-17 | Nicaragua |
| EPI_ISL_17062276 | 2022-11-22 | Nicaragua |
| EPI_ISL_17062277 | 2022-11-24 | Nicaragua |
| EPI_ISL_17062278 | 2022-11-24 | Nicaragua |
| EPI_ISL_17062279 | 2022-11-25 | Nicaragua |
| EPI_ISL_17062280 | 2022-11-25 | Nicaragua |
| EPI_ISL_17062281 | 2022-11-26 | Nicaragua |
| EPI_ISL_17062282 | 2022-11-26 | Nicaragua |
| EPI_ISL_17062283 | 2022-11-27 | Nicaragua |
| EPI_ISL_17062284 | 2022-11-27 | Nicaragua |
| EPI_ISL_17063868 | 2022-11-28 | Nicaragua |
| EPI_ISL_17062285 | 2022-11-28 | Nicaragua |
| EPI_ISL_17062286 | 2022-11-29 | Nicaragua |
| EPI_ISL_17062287 | 2022-11-29 | Nicaragua |
| EPI_ISL_17062288 | 2022-11-29 | Nicaragua |
| EPI_ISL_17062289 | 2022-11-29 | Nicaragua |
| EPI_ISL_17062290 | 2022-11-30 | Nicaragua |
| EPI_ISL_17062291 | 2022-12-02 | Nicaragua |
| EPI_ISL_17062292 | 2022-12-04 | Nicaragua |
| EPI_ISL_17062293 | 2022-12-04 | Nicaragua |
| EPI_ISL_17062294 | 2022-12-04 | Nicaragua |
| EPI_ISL_17062295 | 2022-12-04 | Nicaragua |
| EPI_ISL_17062296 | 2022-12-05 | Nicaragua |
| EPI_ISL_17062297 | 2022-12-05 | Nicaragua |
| EPI_ISL_17062298 | 2022-12-05 | Nicaragua |
| EPI_ISL_17062299 | 2022-12-05 | Nicaragua |
| EPI_ISL_17062300 | 2022-12-05 | Nicaragua |
| EPI_ISL_17062301 | 2022-12-06 | Nicaragua |
| EPI_ISL_17062302 | 2022-12-06 | Nicaragua |
| EPI_ISL_17062303 | 2022-12-06 | Nicaragua |
| EPI_ISL_17062304 | 2022-12-06 | Nicaragua |
| EPI_ISL_17062305 | 2022-12-06 | Nicaragua |
| EPI_ISL_17062306 | 2022-12-06 | Nicaragua |
| EPI_ISL_17062307 | 2022-12-06 | Nicaragua |
| EPI_ISL_17062308 | 2022-12-06 | Nicaragua |
| EPI_ISL_17062309 | 2022-12-06 | Nicaragua |
| EPI_ISL_17062310 | 2022-12-06 | Nicaragua |
| EPI_ISL_17062311 | 2022-12-06 | Nicaragua |
| EPI_ISL_17062312 | 2022-12-06 | Nicaragua |
| EPI_ISL_17062313 | 2022-12-06 | Nicaragua |
| EPI_ISL_17062314 | 2022-12-06 | Nicaragua |
| EPI_ISL_17062315 | 2022-12-06 | Nicaragua |
| EPI_ISL_17062316 | 2022-12-06 | Nicaragua |
| EPI_ISL_17062317 | 2022-12-07 | Nicaragua |
| EPI_ISL_17062318 | 2022-12-07 | Nicaragua |
| EPI_ISL_17062319 | 2022-12-07 | Nicaragua |
| EPI_ISL_17062320 | 2022-12-07 | Nicaragua |
| EPI_ISL_17062321 | 2022-12-07 | Nicaragua |
| EPI_ISL_17062322 | 2022-12-07 | Nicaragua |
| EPI_ISL_17062323 | 2022-12-07 | Nicaragua |
| EPI_ISL_17062324 | 2022-12-07 | Nicaragua |
| EPI_ISL_17062325 | 2022-12-07 | Nicaragua |
| EPI_ISL_17063869 | 2022-12-07 | Nicaragua |
| EPI_ISL_17062326 | 2022-12-07 | Nicaragua |
| EPI_ISL_17062327 | 2022-12-07 | Nicaragua |
| EPI_ISL_17062328 | 2022-12-07 | Nicaragua |
| EPI_ISL_17062329 | 2022-12-07 | Nicaragua |
| EPI_ISL_17062330 | 2022-12-08 | Nicaragua |
| EPI_ISL_17062331 | 2022-12-08 | Nicaragua |
| EPI_ISL_17062332 | 2022-12-08 | Nicaragua |
| EPI_ISL_17062333 | 2022-12-08 | Nicaragua |
| EPI_ISL_17062334 | 2022-12-08 | Nicaragua |
| EPI_ISL_17062335 | 2022-12-08 | Nicaragua |
| EPI_ISL_17062336 | 2022-12-08 | Nicaragua |
| EPI_ISL_17062337 | 2022-12-08 | Nicaragua |
| EPI_ISL_17062338 | 2022-12-08 | Nicaragua |
| EPI_ISL_17062339 | 2022-12-08 | Nicaragua |
| EPI_ISL_17062340 | 2022-12-08 | Nicaragua |
| EPI_ISL_17062341 | 2022-12-08 | Nicaragua |
| EPI_ISL_17062342 | 2022-12-08 | Nicaragua |
| EPI_ISL_17062343 | 2022-12-08 | Nicaragua |
| EPI_ISL_17062344 | 2022-12-09 | Nicaragua |
| EPI_ISL_17062345 | 2022-12-09 | Nicaragua |
| EPI_ISL_17062346 | 2022-12-09 | Nicaragua |
| EPI_ISL_17062347 | 2022-12-09 | Nicaragua |
| EPI_ISL_17062348 | 2022-12-09 | Nicaragua |
| EPI_ISL_17062349 | 2022-12-09 | Nicaragua |
| EPI_ISL_17062350 | 2022-12-09 | Nicaragua |
| EPI_ISL_17062351 | 2022-12-09 | Nicaragua |
| EPI_ISL_17062352 | 2022-12-09 | Nicaragua |
| EPI_ISL_17062353 | 2022-12-09 | Nicaragua |
| EPI_ISL_17062354 | 2022-12-09 | Nicaragua |
| EPI_ISL_17062355 | 2022-12-09 | Nicaragua |
| EPI_ISL_17062356 | 2022-12-10 | Nicaragua |
| EPI_ISL_17062357 | 2022-12-10 | Nicaragua |
| EPI_ISL_17062358 | 2022-12-10 | Nicaragua |
| EPI_ISL_17062359 | 2022-12-10 | Nicaragua |
| EPI_ISL_17062360 | 2022-12-10 | Nicaragua |
| EPI_ISL_17062361 | 2022-12-10 | Nicaragua |
| EPI_ISL_17062362 | 2022-12-10 | Nicaragua |
| EPI_ISL_17062363 | 2022-12-10 | Nicaragua |
| EPI_ISL_17062364 | 2022-12-10 | Nicaragua |
| EPI_ISL_17062365 | 2022-12-10 | Nicaragua |
| EPI_ISL_17062366 | 2022-12-10 | Nicaragua |
| EPI_ISL_17062367 | 2022-12-10 | Nicaragua |
| EPI_ISL_17062368 | 2022-12-11 | Nicaragua |
| EPI_ISL_17062369 | 2022-12-12 | Nicaragua |
| EPI_ISL_17062370 | 2022-12-12 | Nicaragua |
| EPI_ISL_17062371 | 2022-12-12 | Nicaragua |
| EPI_ISL_17062372 | 2022-12-12 | Nicaragua |
| EPI_ISL_17062373 | 2022-12-12 | Nicaragua |
| EPI_ISL_17065719 | 2021-12-31 | Nicaragua |
| EPI_ISL_17065420 | 2022-08-17 | Nicaragua |
| EPI_ISL_17065419 | 2022-08-18 | Nicaragua |
| EPI_ISL_17065426 | 2022-08-15 | Nicaragua |
| EPI_ISL_17065422 | 2022-08-16 | Nicaragua |
| EPI_ISL_17065421 | 2022-08-16 | Nicaragua |
| EPI_ISL_17065733 | 2022-08-25 | Nicaragua |
| EPI_ISL_17065409 | 2022-08-20 | Nicaragua |
| EPI_ISL_17065412 | 2022-08-22 | Nicaragua |
| EPI_ISL_17065414 | 2022-08-24 | Nicaragua |
| EPI_ISL_17065423 | 2022-08-16 | Nicaragua |
| EPI_ISL_17065410 | 2022-08-20 | Nicaragua |
| EPI_ISL_17065418 | 2022-08-25 | Nicaragua |
| EPI_ISL_17065416 | 2022-08-25 | Nicaragua |
| EPI_ISL_17065425 | 2022-08-15 | Nicaragua |
| EPI_ISL_17065424 | 2022-08-15 | Nicaragua |
| EPI_ISL_17065415 | 2022-08-24 | Nicaragua |
| EPI_ISL_15326870 | 2021-06-10 | Nicaragua |
| EPI_ISL_17065544 | 2021-10-07 | Nicaragua |
| EPI_ISL_17065545 | 2021-10-07 | Nicaragua |
| EPI_ISL_17065546 | 2021-10-07 | Nicaragua |
| EPI_ISL_17065547 | 2021-10-09 | Nicaragua |
| EPI_ISL_17065548 | 2021-10-09 | Nicaragua |
| EPI_ISL_17065542 | 2021-10-10 | Nicaragua |
| EPI_ISL_17065550 | 2021-10-10 | Nicaragua |
| EPI_ISL_17065549 | 2021-10-10 | Nicaragua |
| EPI_ISL_17065551 | 2021-10-11 | Nicaragua |
| EPI_ISL_17065543 | 2021-10-26 | Nicaragua |
| EPI_ISL_17065553 | 2021-10-27 | Nicaragua |
| EPI_ISL_17065552 | 2021-10-27 | Nicaragua |
| EPI_ISL_17065554 | 2021-11-16 | Nicaragua |
| EPI_ISL_17065849 | 2022-01-29 | Nicaragua |
| EPI_ISL_17065413 | 2022-08-24 | Nicaragua |
| EPI_ISL_17065411 | 2022-08-22 | Nicaragua |
| EPI_ISL_17065417 | 2022-08-25 | Nicaragua |
| EPI_ISL_17065761 | 2021-10-09 | Nicaragua |
| EPI_ISL_17065762 | 2021-10-07 | Nicaragua |
| EPI_ISL_17065763 | 2021-10-09 | Nicaragua |
| EPI_ISL_17065764 | 2021-10-11 | Nicaragua |
| EPI_ISL_17065765 | 2021-10-11 | Nicaragua |
| EPI_ISL_17065766 | 2021-10-11 | Nicaragua |
| EPI_ISL_17065767 | 2021-10-21 | Nicaragua |
| EPI_ISL_17065768 | 2021-10-23 | Nicaragua |
| EPI_ISL_17065769 | 2021-10-27 | Nicaragua |
